# Dosage sensitivity in Pumilio1-related phenotypes reflects distinct disease mechanisms

**DOI:** 10.1101/2021.03.11.435015

**Authors:** Salvatore Botta, Nicola de Prisco, Alexei Chemiakine, Maximilian Cabaj, Purvi Patel, Ella Doron-Mandel, Colton J. Treadway, Marko Jovanovic, Nicholas G. Brown, Rajesh K. Soni, Vincenzo A. Gennarino

## Abstract

Mutations in the RNA-binding protein (RBP) Pumilio1 (PUM1) can cause dramatically different phenotypes. We previously noted that phenotypic severity tracked with protein dosage: a mild mutation that reduces PUM1 levels by 25% causes late-onset ataxia, whereas PUM1 haploinsufficiency causes developmental delay and seizures. Why this difference in expression should cause such different phenotypes has been unclear: PUM1 targets are de-repressed to equal degrees in both cases, and the more severe mutation does not hinder PUM1’s RNA-binding ability. We therefore developed a PUM1 interactome in the murine brain. We find that mild PUM1 loss de-represses PUM1-specific targets, but PUM1 haploinsufficiency disrupts several interactors and regulation of their targets. We validated these phenomena in patient-derived cell lines and show that normalizing PUM1 levels restores interactors and their targets to proper levels. We therefore propose that dosage sensitivity does not necessarily reflect a linear change in protein abundance but can involve distinct mechanisms. Studying the interactors of RBPs *in vivo* will be necessary to understand their functions in neurological diseases.

## Introduction

RNA-binding proteins (RBPs) modify the proteome at the post-transcriptional level, regulating RNA localization, transport, translation, splicing, and decay; they have been found to orchestrate hundreds of pathways that are responsible for proper biological functions (Hentze *et al*, 2018; Keene, 2007b). As our understanding of RBPs has grown over the past decade, it has become clear that their functions are particularly important in neurons, whose synaptic plasticity demands rapid local responses to stimuli (Mauger *et al*, 2016). Several complex neurological disorders—e.g., Fragile X syndrome, Fragile X-associated tremor and ataxia syndrome, and amyotrophic lateral sclerosis—are known to involve disruptions in the function of RBPs, which has understandably led to considerable interest in mapping their neuronal targets (Darnell & Richter, 2012; Khalil *et al*, 2018; Ravanidis *et al*, 2018).

Our work on Pumilio1 (PUM1), however, has revealed that the targets may not provide a full picture of the RBP’s functioning. After discovering that PUM1 is important for mouse neurobiology (Gennarino *et al*, 2015), we searched human databases for individuals with mutations in *PUM1*. We initially found 13 individuals with deletions or missense mutations, and their phenotypes tracked with protein dosage. A mild mutation (T1035S) that reduces PUM1 levels by 25% causes a slowly progressive, pure ataxia with onset in mid-life, which we called PUM1-related cerebellar ataxia or PRCA. Deletions or the most severe missense mutation (R1147W) reduce PUM1 levels by ∼50% and cause a neurodevelopmental syndrome we called PUM1-associated developmental delay and seizures, or PADDAS (Gennarino *et al*, 2018). Both the infantile and adult-onset phenotypes are now considered to be forms of spinocerebellar ataxia type 47 (SCA47). T1035S lies within a highly conserved RNA-binding domain and impairs RNA binding, so it was not surprising to find known PUM1 targets upregulated in patient cells. The puzzle came with R1147W, which lies outside this domain and does not impair target regulation, yet deregulates the same targets to the same degree as T1035S (Gennarino *et al*., 2018). Moreover, the R1147W phenotype is quite severe, closer to that of the null mice than the heterozygous mice. Given that RBPs can interact with, influence, or compete with each other (Dassi, 2017), we hypothesized that R1147W disrupts PUM1’s ability to interact with its native partners, which would then lead to de-repression of the targets of these complexes and not just the direct targets of PUM1.

Testing this hypothesis required us to identify PUM1 interactors in the mouse brain. Although a great deal is known about the PUMILIO/FBF (PUF) family of RBPs (Bohn *et al*, 2018; Friend *et al*, 2012; Goldstrohm *et al*, 2018; Kedde *et al*, 2010; Miles *et al*, 2012; Temme *et al*, 2014; Uyhazi *et al*, 2020; Van Etten *et al*, 2012; Weidmann *et al*, 2014), of which PUM1 and its homolog PUM2 are members, little is known about PUM1 function in the postnatal mammalian brain. Protein interactions in general, and those of PUF (Pumilio and FBF) family members in particular, can be organism-, transcript-, and even condition-specific (Marrero *et al*, 2011). We therefore took an unbiased approach by using *in vivo* proteomics to identify PUM1’s native partners in the mouse brain. We then studied the effect of PUM1 insufficiency on a subset of RBP interactors in *PUM1* heterozygous and null mice and cell lines from patients bearing either the T1035S or R1147W mutation. The results support our hypothesis that two different mechanisms underlie PRCA and PADDAS, underscoring the need to examine an RBP’s interactions as well as identify its targets.

## Results

### Establishing the PUM1 interactome in the adult mouse brain

We performed co-immunoprecipitation (IP) on brains from 10-week-old wild-type (WT) mice followed by liquid chromatography with tandem mass spectrometry (LC-MS/MS); we used IgG as a negative control (Figure S1A). Principal component analysis (PCA) separated IP-IgG from IP-Pum1 (Figure S1B). No residual Pum1 was detected in brain tissue by post-IP western blot (Figure S1C). (In this paper, we will differentiate between human and mouse proteins by using all-caps only for the former.) Considering only those candidates that had at least two unique peptides in at least five out of six IP-Pum1 samples, this approach identified 234 putative interactors (Figure S1C and **Table S1**) that we clustered into 10 functional groups using protein-protein interaction data from CORUM and the Human Protein Atlas (Raudvere *et al*, 2019) (Figure 1).

**Figure 1.**
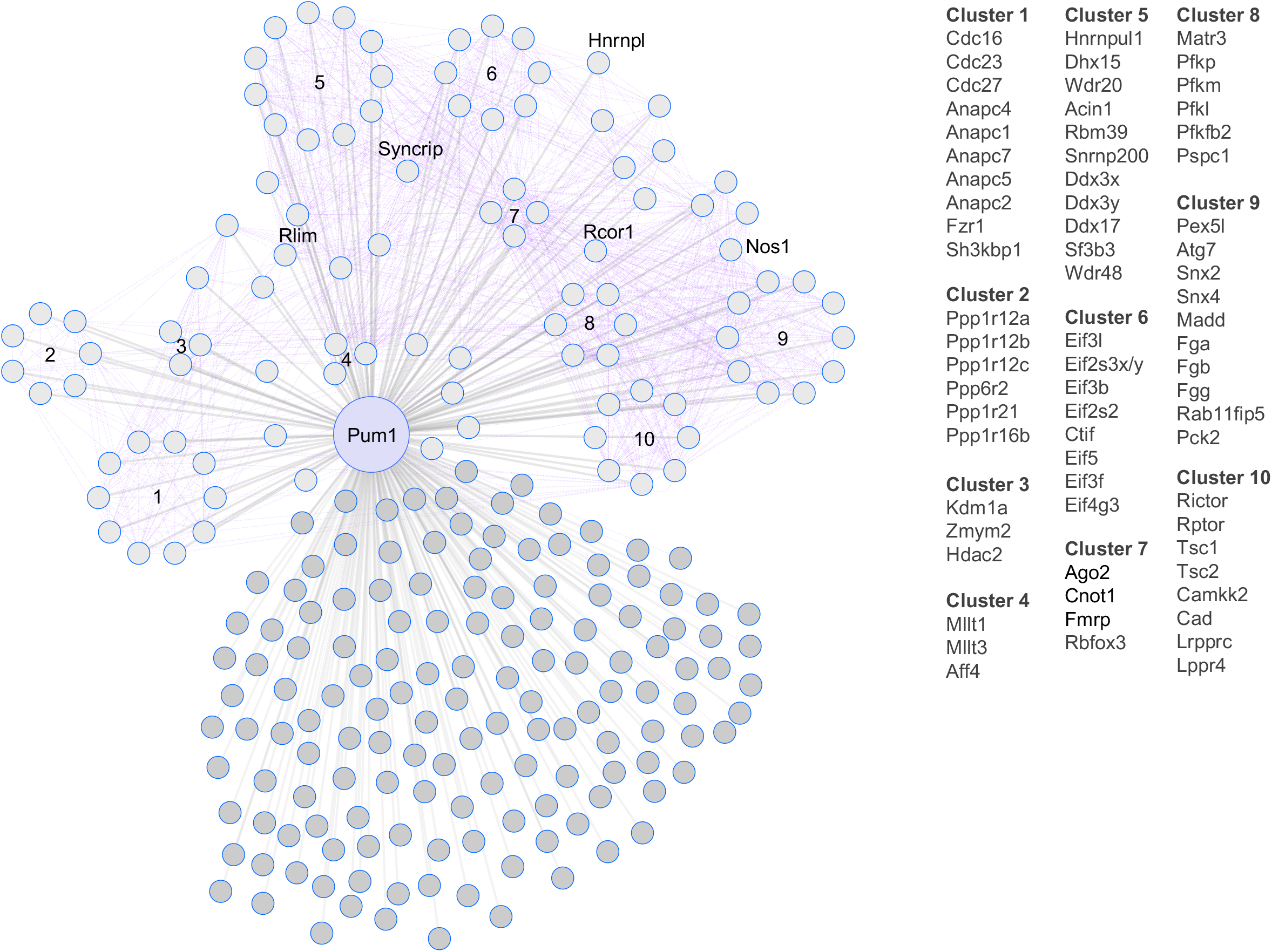
A brain-specific Pum1 interactome. Network of putative Pum1 interactors in 10-week-old mouse brain (circles connected to Pum1 by gray lines). Interactions between interactors (purple lines) were inferred by g:GOSt from Corum and the Human Protein Atlas (see Materials and Methods). The proteins in each of the 10 clusters are listed to the right. We combined and homogenized whole brains from two 10-week-old wild-type mice per sample (1 female and 1 male), aliquoting half of each sample for IP against either Pum1 or IgG, then performed six biological replicates (six samples, 12 mice total) for each LC-MS/MS experiment against IP-Pum1 and IP-IgG. All putative Pum1 interactors are listed in Table S1.

Our list included several proteins that had been previously found to interact with Pum1, as well as new members of protein complexes or families that interact with Pum1 in other contexts. For example, our LC-MS/MS identified Fmrp (cluster 7), which associates with Pum1 in neural progenitor cells (Zhang *et al*, 2017). We also identified Cnot1, the central scaffold of the CCR4-NOT complex (Enwerem *et al*, 2021; Van Etten *et al*., 2012), which is recruited by Pum1 to shorten poly(A) tails and promote mRNA degradation (Temme *et al*., 2014; Van Etten *et al*., 2012; Weidmann *et al*., 2014). Translation initiation factors (cluster 6) cooperate with PUF proteins in invertebrates (Blewett & Goldstrohm, 2012).

*Drosophila* studies showed that Pumilio partners with different proteins in different neuronal types (Muraro *et al*, 2008), so we repeated the LC-MS/MS experiments in three brain regions that express abundant Pum1: the cerebellum, hippocampus, and cortex (Gennarino *et al*., 2015). PCA readily separated Pum1 and IgG samples (Figure S2A). This analysis identified 854 putative Pum1 interactors in the cerebellum, 423 in the hippocampus, and 598 in the cortex (Figure 2A, Figure S2B-D and **Table S1**). 489 were unique to the cerebellum, 145 to the hippocampus, 247 to the cortex, and 48 unique to the rest of the brain (i.e., excluding these three regions). Only 82 candidates appeared in all three brain regions and the whole brain (Figure 2A*, yellow dots*).

**Figure 2.**
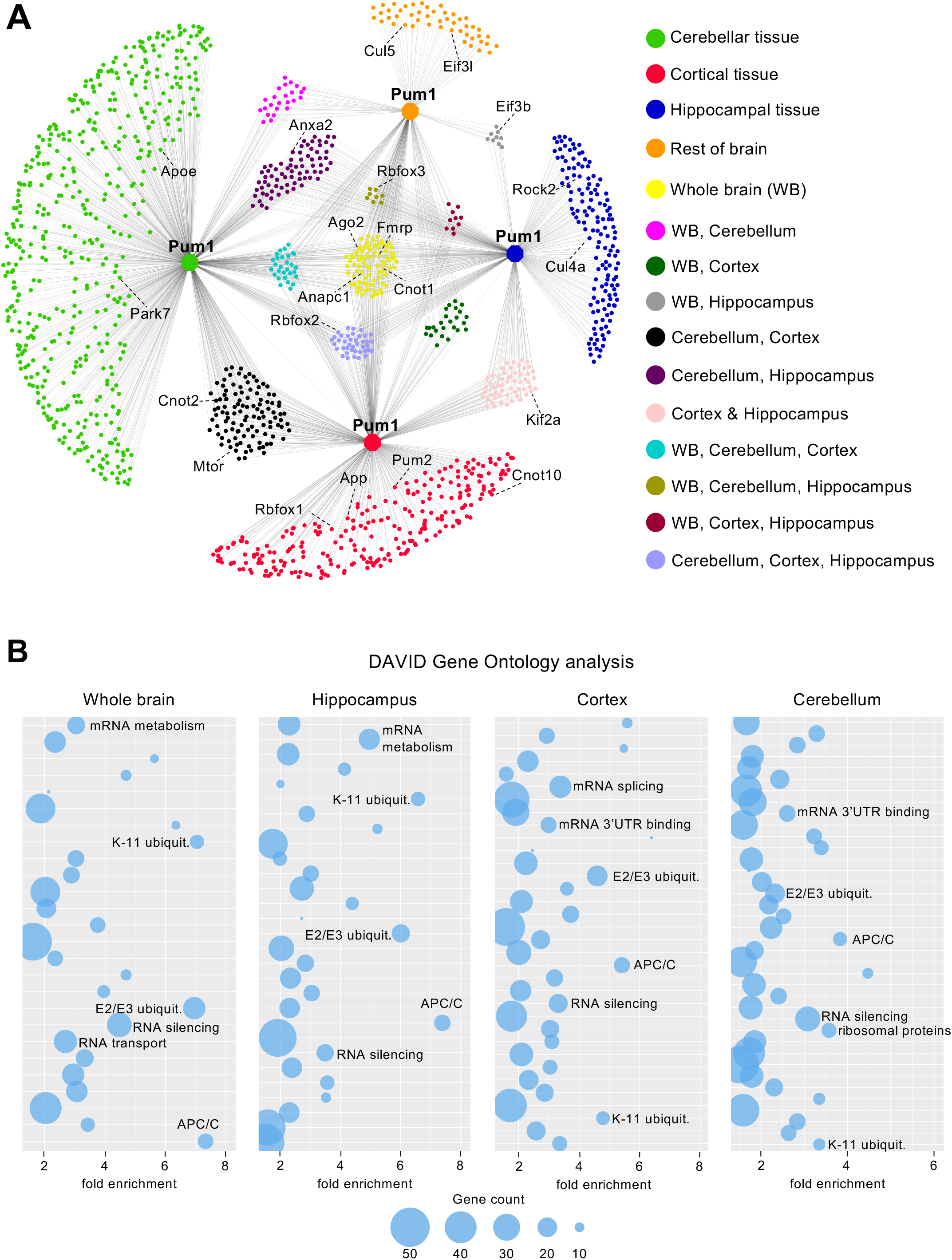
A brain-region specific Pum1 interactome. (**A**) Pum1 interactome from 10-week-old mouse cerebellum (n=8 mice, 4 male and 4 female), hippocampus (n=10, 5/5), cortex (n=8, 4/4) and the rest of the brain (i.e., excluding those three regions) for a total of 1,500 proteins (**Table S1**). Node colors represent different brain regions or the overlap between two or more brain regions as noted. All experiments were performed at least in triplicate. IP against IgG was used as a negative control. (**B**) Bubble plots show the top categories from gene ontology analyses of Pum1 interactors from whole brain, hippocampus, cortex, and cerebellum. Only the categories with fold enrichment >1.5 and FDR<0.05 are shown; not all are labeled because of space limitations. The full list of gene ontology categories is available in **Table S2**.

To determine whether these results merely reflect regional differences in protein abundance, we performed quantitative proteomics in 10-week-old whole brain, cerebellum, cortex, and hippocampus. Pum1 interactors in general were not the most abundant proteins in these samples (Figure S3A, B). For example, Pum2, which turned up in only 3 out of 6 of the whole-brain samples, was one of the strongest Pum1 interactors in the cortex (Figure 2A, S3C, and **Table S1**). It is expressed at low levels in all brain regions examined (Figure S3D). These findings suggest that the candidate interactors represent biological information rather than post-lysis effects or high abundance.

Components of the APC/C and mTOR pathways were consistent across the three brain regions (Figure 2A, B, and **Table S1**), but the list of Pum1 interactors in several other pathways was expanded by this regional analysis (Figure 2). For example, although Cnot1 turned up in all three brain regions (Figure S3C), Cnot2 appeared only in cortex and cerebellum, and Cnot10 only in the cortex (Figure 2A). There were many putative interactors involved in translation initiation, with Eif3b showing up in both the hippocampus and the whole brain (Figure 2A). The splicing factors showed regional specificity in interactions: Rbfox1 was restricted to the cortex, while Rbfox2 and Rbfox3 were in the cerebellum, hippocampus, and whole brain (Figure 2A). This is consistent with previous work showing that Rbfox1 mediates cell-type-specific splicing in cortical interneurons (Wamsley *et al*, 2018) and that Rbfox2 is needed for cerebellar development (Gehman *et al*, 2012).

DAVID Gene Ontology analysis revealed that the main functional categories across the three regions were ubiquitin ligases (anaphase-promoting complex [APC/C], E2/E3 and Kll-linked ubiquitin) and RBPs involved in various aspects of RNA metabolism (RNA silencing, 3’UTR binding, mRNA stability, transport, and splicing) (Figure 2B and **Table S2**). To test some of these interactions biochemically, we tried to purify recombinant proteins, but many of the proteins or their complexes are either quite large (>100kD) or strongly tended to aggregate. We were able pull down PUM1 by recombinant GST-tagged FMRP (Figure S4A) and GST-tagged PUM2 (Figure S4B), confirming their direct interactions (Figure S4). We also used IP and co-IP to spot-check key candidates from the anaphase promoting complex or cyclosome (APC/C, cluster 1: Anapc1, Fzr1) and mTOR (cluster 10: Rptor, Cad) pathways (Figure S3C, S4C, and D).

Realizing we would need to select certain proteins for further *in vivo* study, we decided to prioritize the RBPs in cluster 7 (Figure 1): Fmrp and Ago2 (involved in RNA silencing), Cnot1 (mRNA deadenylase protein), Rbfox3 (alternative splicing factor), and Pum2. We reasoned as follows. First, RBP-RBP interactions are biologically important, and RNA-related categories were prominent in the gene ontology analyses for both whole brain and all three brain regions. Second, this cluster was also the most highly interconnected with other clusters and likely to influence their activities. Third, these RBPs have been well studied, which would enable us to more readily test the consequences of PUM1 loss. Nevertheless, these proteins have been studied mostly *in vitro* and, with the exception of Fmrp and Pum2, have not been previously associated with Pum1 in the murine brain, so they might shed more light on Pum1 biology.

### Pum1 associates with Fmrp, Ago2, Cnot1, and Pum2 in the absence of mRNA

We confirmed Pum1 associations with Pum2, Fmrp, Ago2, Rbfox3, and Cnot1 by co-IP followed by western blot (Figure 3A, *left panel*). We also tested for murine Mov10, which binds both Fmrp and Ago2 *in vitro* (Kenny *et al*, 2020; Kenny *et al*, 2014), and found it pulled down with Pum1, likely in concert with Fmrp (Figure 3A). We co-IP’d Pum1 and blotted for all six RBPs in *Pum1^-/-^* mouse brains; none were detected (Figure S5A). We then tested other proteins associated with the RNA silencing machinery that did not appear in our LC-MS/MS data, such as Ago1 and Ago3, and we found no interactions (Figure S5B).

**Figure 3.**
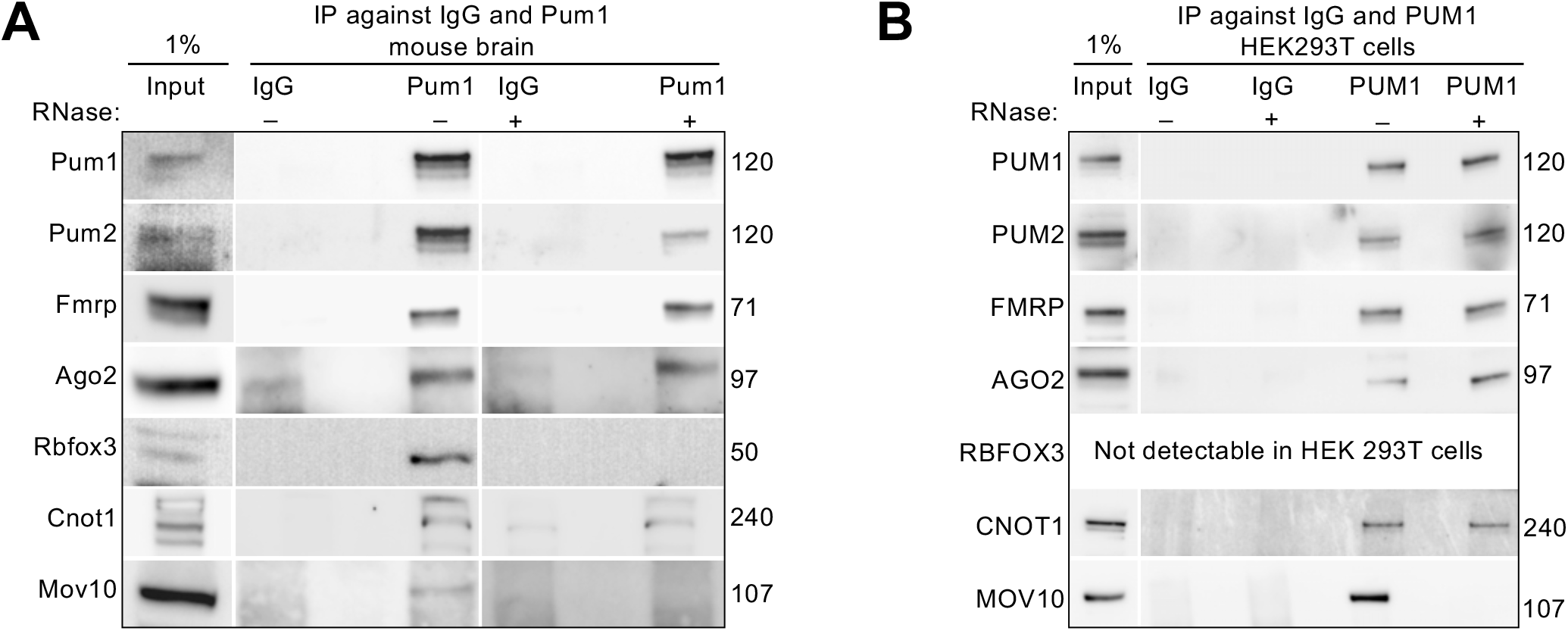
Validation of Pum1 associations with post-transcriptional RNA-binding proteins in mouse brain and HEK293T cells. (**A**) Representative western blot of proteins pulled down by IP against Pum1 compared to IgG from wild-type mice brain without (*left*) and with (*right*) RNase treatment. In this panel, after IP-Pum1, we immunoblotted for Pum1 (positive control), Pum2, Fmrp, Ago2, Rbfox3, Cnot1, and Mov10. (see Materials and Methods.). (**B**) Representative western blots of the same proteins validated in panels **A** after IP against PUM1 with or without RNase treatment from HEK293T cell lines. In panels **A** and **B**, IP against IgG was used as a negative control and Input (1% from the initial protein lysate) as a loading control. The numbers on the right are the respective molecular weights expressed in kilodaltons (kDa). All the experiments were repeated at least four times. All mice were sacrificed at 10 weeks of age.

To exclude the possibility that the co-IP experiments were recovering proteins that co-bound target RNAs but are not part of the same complex as the protein of interest, we treated mouse brain samples with RNase and verified that no detectable RNA remained (see Materials and Methods). Pum1 still associated with Pum2, Fmrp, Ago2, and Cnot1 in the absence of mRNA, but not with Rbfox3 or Mov10 (Figure 3A, *right panel*, and **S5C**). We repeated the RNase experiments in HEK293T cells, which confirmed our results (except for Rbfox3, which was not detectable in these cells) (Figure 3B). These data suggest that Pum1 interacts with Pum2, Fmrp, Ago2, and Cnot1 prior to binding RNA.

### Pum1 loss alters RBP interactors and miRNA machinery in mouse cerebella by sex

If Pum1 is an important interactor for these six RBPs, loss of Pum1 should affect their stability or abundance in mouse brain. *Pum1* heterozygous and null mice showed changes in the quantities of Pum2, Ago2, and Mov10 proteins across the brain (Figure S6A), but only *Pum2* showed changes in mRNA levels (Figure S6B). Ago2 and Mov10 levels fell only in male mice (Figure S6A), which led us to quantify Pum2, Fmrp, Ago2, Rbfox3, Cnot1 and Mov10 mRNA and protein in the cerebellum, cortex, and hippocampus of male and female mice.

As previously reported, Fmrp protein expression was upregulated in all three brain regions in male null mice, but in female null cerebella Fmrp was almost 70% lower (Figure 4A) (Singh *et al*, 2007; Singh & Prasad, 2008). Pum2 protein levels rose in all three brain regions, as did its mRNA (Figure 4A-C, and Figure S7). Ago2, Rbfox3, and Cnot1 also showed divergent responses to Pum1 loss according to sex and brain region (Figure 4A-C). None of these proteins showed any changes in mRNA levels (Figure S7), despite the fact that *Fmr1* and *Cnot1*, like *Pum2*, have a Pumilio Response Element (PRE) (Zamore *et al*, 1997) in their 3’UTR.

**Figure 4.**
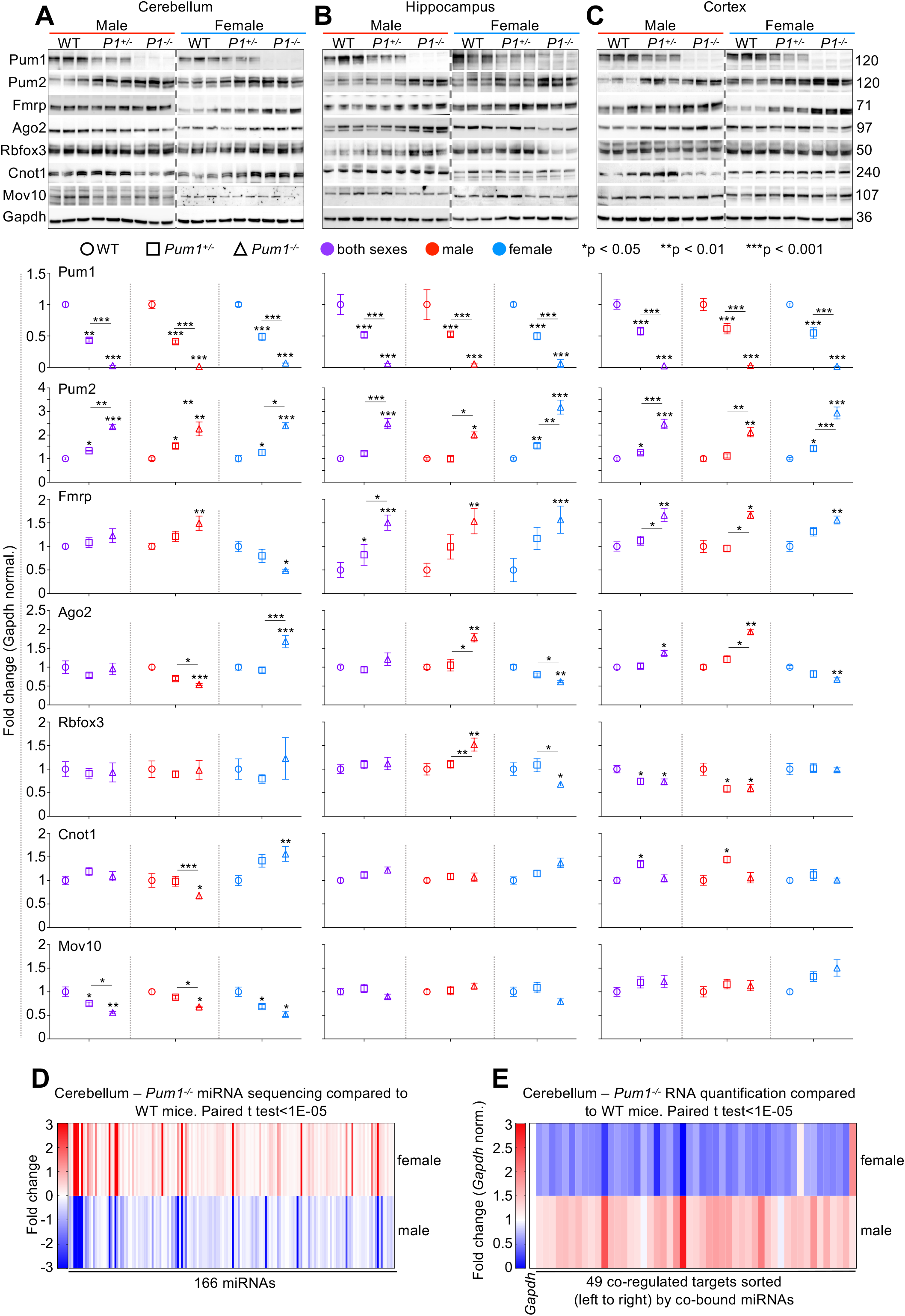
Pum1 interactors and the microRNA machinery show brain region- and sex-specific responses to Pum1 loss. Representative western blots of Pum1, Pum2, Fmrp, Ago2, Rbfox3, Cnot1, and Mov10 in (**A**) cerebellum, (**B**) hippocampus, and (**C**) cortex in both male (left panel) and female (right panel) WT, *Pum1^+/-^*, and *Pum1^-/-^* mice. All the experiments were conducted with equal numbers of 10-week-old male and female mice per genotype, for a total of at least 12 mice per genotype (data represent mean ± SEM). Graphs below show quantification for each protein by brain region, sex, and genotype. All data were normalized to Gapdh protein levels. Data represent mean ± SEM. P values were calculated by two-tailed Student’s t test. \**p* < 0.05, \*\**p* < 0.01, \*\*\**p* < 0.001. P1 indicates Pum1. See Figure S8 for mRNA quantification for each interactor, brain region, and sex. (**D**) Heatmap showing 166 microRNAs from cerebella of *Pum1^-/-^*male and female mice that were dysregulated (fold change −3 to +3) relative to wild-type cerebellum. The full list of miRNA names and fold changes are available in Table S3. See Figure S9 for male and female miRNA scatter plots. (**E**) Heatmap showing mRNA quantification by qPCR for 49 targets co-bound by a minimum of eight dysregulated miRNAs (>25% change) from panel **D**. Statistical significance and magnitude of dysregulation are illustrated for both male and female in Figure S10. The entire list of targets predicted to be co-bound by at least two miRNAs is presented in Table S4.

Because Pum1 loss has deleterious effects on the cerebellum in both mice (Gennarino *et al*., 2015) and humans (Gennarino *et al*., 2018), and Pum1 interacts closely with the miRNA machinery (Kedde *et al*., 2010), we decided to investigate the physiological relevance of these sex-specific observations by examining the effects of divergent Ago2 protein levels on cerebellar miRNAs in male and female mice. Our miRNA-seq identified 701 expressed miRNAs, many of which diverged in expression between the two sexes (Figure S8A, B). Heatmap clustering of significant miRNA expression in *Pum1^-/-^* and WT male and female cerebella at 10 weeks of age revealed that the expression of 166 miRNAs diverged between the two sexes in parallel with Ago2 expression (**Table S3,** Figure 4D).

To study the expression of downstream targets that are co-bound by those miRNAs in 10-week-old WT and *Pum1^-/-^* male and female cerebella, we selected the 49 miRNAs that showed >25% change in expression in either direction. Using TargetScan and CoMeTa (Agarwal *et al*, 2015; Gennarino *et al*, 2012) we identified 6832 putative targets that are co-bound by at least two miRNA. To reduce this list to manageable size, we prioritized targets that are co-bound by at least eight miRNAs, for a total of, coincidentally, 49 putative targets. *Pum1^-/-^* male and female cerebella showed gene expression changes for 44 out of these 49 targets that correlated with the sex differences in Ago2 levels (Figure 4E, Figure S9 and **Table S4**).

To elucidate the biological pathways in which these miRNAs play an essential role, we performed David Gene Ontology (GO) with all the non-redundant targets predicted by CoMeTa (Gennarino *et al*., 2012) and TargetScan 7.1 (Agarwal *et al*., 2015) that are co-bound by at least four miRNAs (GO needs a larger pool of genes than 49 but fewer than 3000), 2127 targets (Figure S10A-C). Among 2127 targets analyzed, we found enrichment for multiple categories having to do with synaptic function under “cellular components.” The most enriched categories under “biological processes” were organ growth and post-embryonic development. Under KEGG pathways, there was a particular enrichment in Wnt signaling, dopaminergic and cholinergic pathways, cancers, and protein ubiquitination.

We then analyzed the same targets by SynGO (Koopmans *et al*, 2019), which uses single-cell data to identify genes that are expressed in specific neurons. SynGO pinpointed 117 presynpatic targets and 124 postsynaptic (Figure S10D). Among the 166 miRNAs inversely expressed between sexes were the entire miR-200 family (miR-200a, miR-220b, miR-200c, miR-141, and miR-429), which regulates targets involved in neurogenesis, glioma, and neurodegenerative diseases (Fu *et al*, 2019; Trumbach & Prakash, 2015). These results are consonant with SCA47 cerebellar deficits and suggest an intimate relation between Pum1 and Ago2 in mouse cerebellum.

### Pum1, Pum2, Fmrp, Ago2, and Rbfox3 share their top targets

If the complexes Pum1 forms with these RBPs are physiologically relevant, as seen for Ago2 in cerebellum, then they should co-regulate at least some of the same mRNA targets. Indeed, one corollary of the “regulon theory,” which posits that mRNA targets in the same pathway are co-regulated (Blackinton & Keene, 2014; Keene, 2007a, b; Keene & Lager, 2005), is that there should be a discernible set of RBPs that do the co-regulating.

To test this hypothesis, we analyzed all the available high-throughput sequencing UV-crosslinking and immunoprecipitation (HITS-CLIP) data available for the murine brain. These data exist for Fmrp (Maurin *et al*, 2018), Ago2 (Chi *et al*, 2009), Rbfox3 (Weyn-Vanhentenryck *et al*, 2014), Pum1, and Pum2 (Zhang *et al*., 2017)). We then performed gene set enrichment analysis (GSEA) (Subramanian *et al*, 2005) using Fmrp as the basis for comparison (because it has the largest dataset). As negative controls, we used HITS-CLIP data from mouse brain for the other four available RBPs that did not show up as Pum1 interactors in our LC-MS/MS: Mbnl2 (Charizanis *et al*, 2012), Nova (Zhang *et al*, 2010), Apc (Preitner *et al*, 2014), and Ptpb2 (Licatalosi *et al*, 2012).

Although the datasets are not perfectly matched (the mice were different ages in some cases; see Methods), this analysis nonetheless revealed that Pum1 targets were concentrated in the top 5th percentile of all Fmrp targets, with an enrichment score (ES) of 0.93 (the maximum is 1) and FDR of 0 (Figure 5A, *blue line represents ES*). Pum2, Ago2, and Rbfox3 showed nearly identical patterns. There was no significant overlap between the targets of Fmrp and those of any negative control (Nova had the highest ES, but this was only 0.36 with a rank max of 45^th^ percentile and FDR=1; Figure 5B). Neither Pum1 nor its partner RBP targets were enriched among any of the ranked target lists of the negative controls (Figure S11A).

**Figure 5.**
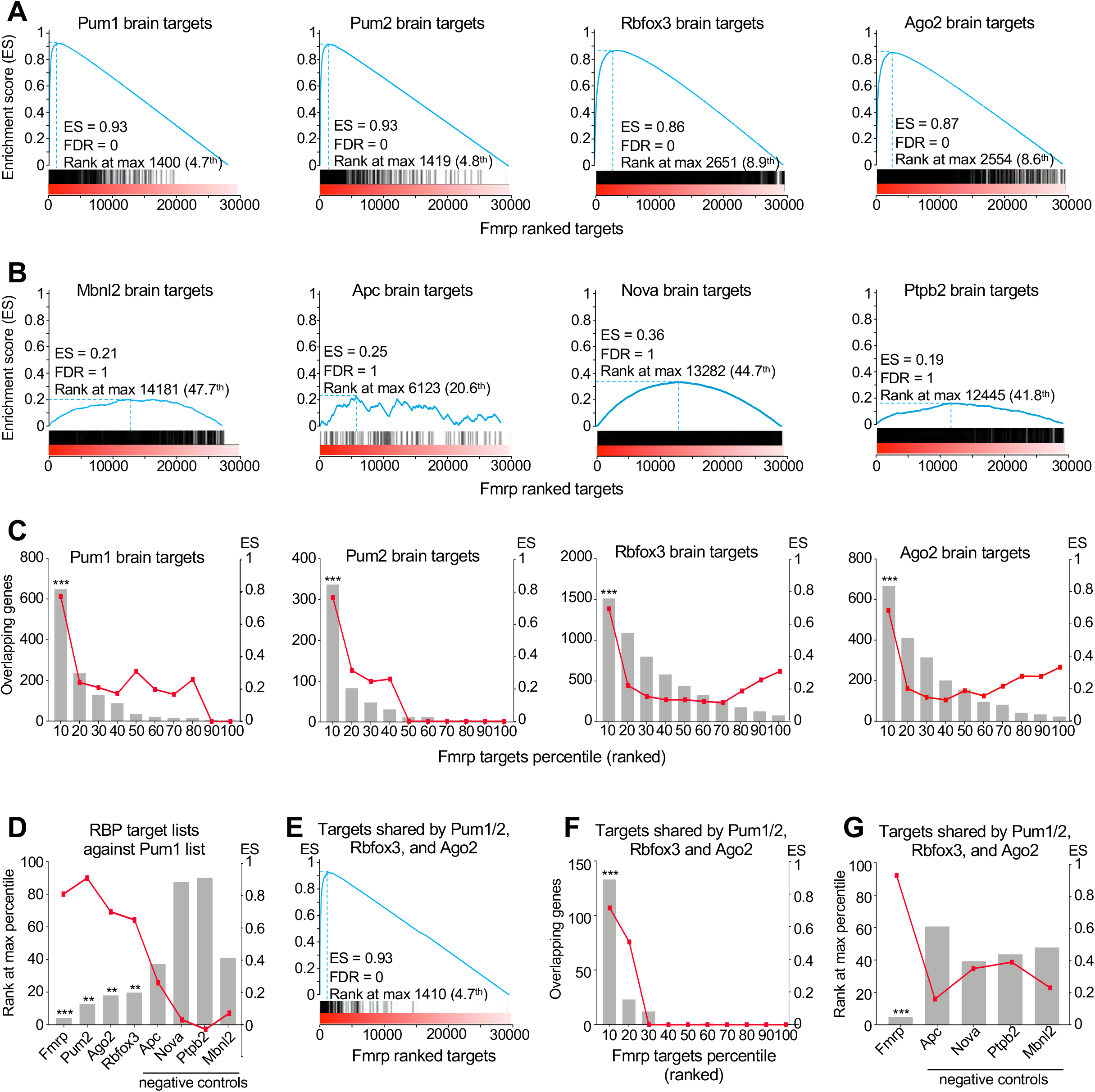
Pum1 and its RNA-binding protein interactors share many neuronal mRNA targets. (**A**) Enrichment plots generated by Gene Set Enrichment Analysis (GSEA) of Pum1, Pum2, Rbfox3, and Ago2 HITS-CLIP targets plotted again Fmrp ranked HITS-CLIP data. Pum1, Pum2, Rbfox3, and Ago2 targets are enriched at the top 10^th^ percentile of the Fmrp targets with FDR=0. (**B**) GSEA analysis scores of HITS-CLIP data from each negative control (Apc, Nova, Ptpb2, and Mbnl2) plotted against Fmrp ranked HITS-CLIP data. The negative controls have a maximum enrichment score of 0.36 for Apc ranking at the top 44.7% with FDR=1. (**C**) GSEA analysis scores of Pum1, Pum2, Rbfox3, and Ago2 HITS-CLIP data plotted against Fmrp HITS-CLIP data divided into 10-percentile ranked bins shows the shared targets are among the top percentiles of targets for each protein. (**D**) GSEA analysis scores of the HITS-CLIP data for Fmrp, Pum2, Ago2, Rbfox3 and four negative controls (Apc, Nova, Ptpb2, and Mbnl2) against Pum1 ranked HITS-CLIP data. The targets of Fmrp, Ago2, Pum2, and Rbfox3 are enriched at the top 5^th^ to 18^th^ percentile of Pum1 targets. (**E**) GSEA analysis of the shared targets between Pum1, Pum2, Ago2, and Rbfox3 against Fmrp showing that they are enriched in the top 5^th^ percentile of Fmrp ranked targets. (**F**) Pum1, Pum2, Ago2, and Rbfox3 shared targets plotted against Fmrp ranked HITS-CLIP targets and divided into 10-percentile bins shows that all of their respective targets are enriched at the top 10^th^ percentile of the Fmrp ranked targets. (**G**) GSEA analysis scores of the targets shared by Pum1, Pum2, Ago2, and Rbfox3 and the four negative controls (Apc, Nova, Ptpb2, and Mbnl2) plotted against Fmrp. At best the negative controls are enriched at the top 40% with a maximum ES of 0.41. For all the GSEA analyses, the False Discovery Rate (FDR) was provided by GSEA: **FDR<0.05 and ***FDR<0.01. ES=Enrichment score (blue line). Note that lowest rank at max percentage indicates stronger targets in the rank (see Materials and Methods). HITS-CLIP data, and the respective rank, were obtained from the literature and were initially acquired as follows: Pum1 and Pum2 (Zhang *et al*., 2017), Fmrp (Maurin *et al*., 2018), Ago2 (Chi *et al*., 2009), Rbfox3 (Weyn-Vanhentenryck *et al*., 2014), Nova (Zhang *et al*., 2010), Ptpb2 (Licatalosi *et al*., 2012), Mbnl2 (Charizanis *et al*., 2012), and Apc (Preitner *et al*., 2014) (see Materials and Methods for more details). The full list of shared targets is reported in Table S5.

To ascertain where highest-ranking Fmrp targets fell among the genes with the highest probability of being Pum1 targets, we divided the Fmrp ranked target list into 10 equal bins according to percentile. We then repeated GSEA of Pum1 HITS-CLIP data for each bin and found that 648 of the 1194 identified Pum1 targets (54%) are in the top 10^th^ percentile of Fmrp targets, with an ES of 0.8 (Figure 5C). This was also true for Pum2, Ago2, and Rbfox3 (Figure 5C).

We then performed the same analysis but instead used the Pum1 target list as the basis for comparison. We ran GSEA on each of the four Pum1 partners against the list of Pum1 target genes, and each partner’s targets were within the top 20% of the Pum1 list (Figure 5D). Specifically, Fmrp’s targets reside in the top 10^th^ percentile (with an ES of 0.81), Pum2’s targets within the 16^th^ percentile (ES=0.9), Ago2’s targets within the 18^th^ percentile (ES=0.76), and Rbofx3’s targets within the 19^th^ percentile (ES=0.67). The four RBPs used here as negative controls have a minimum rank at the 37^th^ percentile, and the best ES was 0.26 for Apc; none of the five reached statistical significance (Figure 5D). These analyses demonstrate that there is substantial overlap among the highest-ranked targets of Pum1, Pum2, Fmrp, Ago2, and Rbfox3.

We also studied the targets shared by Pum1, Pum2, Ago2, and Rbfox3 to determine how they distribute within Fmrp. We found an ES of 0.93 falling within the top 5^th^ percentile (Figure 5E); 141 out of 175 common targets were within the top 10^th^ percentile (bin1) of Fmrp targets, with 99 within the top 5^th^ (Figure 5F). This contrasts with the negative controls, for which the best ES was 0.41 within the top 40^th^-60^th^ percentile (Figure 5G). DAVID gene ontology analysis of those 175 common targets between Ago2, Pum1, Pum2, Fmrp, and Rbfox3 revealed pathways enriched in neurons and axonal projections (Figure S11B, C). Previous studies have shown that Pum1 and Pum2 cooperate with the miRNA machinery to suppress certain targets (Gennarino *et al*., 2015; Kedde *et al*., 2010). Among Fmrp HITS-CLIP targets, there were almost 300 microRNAs. Pum1 HITS-CLIP had 60 miRNAs, only four of which were not shared with Fmrp; Pum2 HITS-CLIP had no miRNAs that were not shared with either Pum1 or Fmrp (Figure S11D and **Table S6**). The Pum1 complexes with these fellow RBPs are therefore physiologically relevant.

### PUM1 interactors are destabilized in cell lines from infantile-onset but not adult-onset patients

Having identified Pum1 interactors and shared targets, we were finally in a position to determine whether either the mildest (T1035S) or most severe (R1147W) missense mutations in PUM1 destabilize these key RBP interactors. We compared patient-derived cell lines (lymphoblasts for T1035S, fibroblasts for R1147W) with their respective age-, sex-, and cell type matched healthy controls. IP (Figure 6A, B) and post-IP (Figure S12A, B) against PUM1 followed by western blot show that our protocol pulled down 100% of PUM1 protein from both patient-derived cell lines. We then confirmed that R1147W, which lies outside the RNA-binding domain, retains RNA-binding activity, while T1035S does not (Figure S12C).

**Figure 6.**
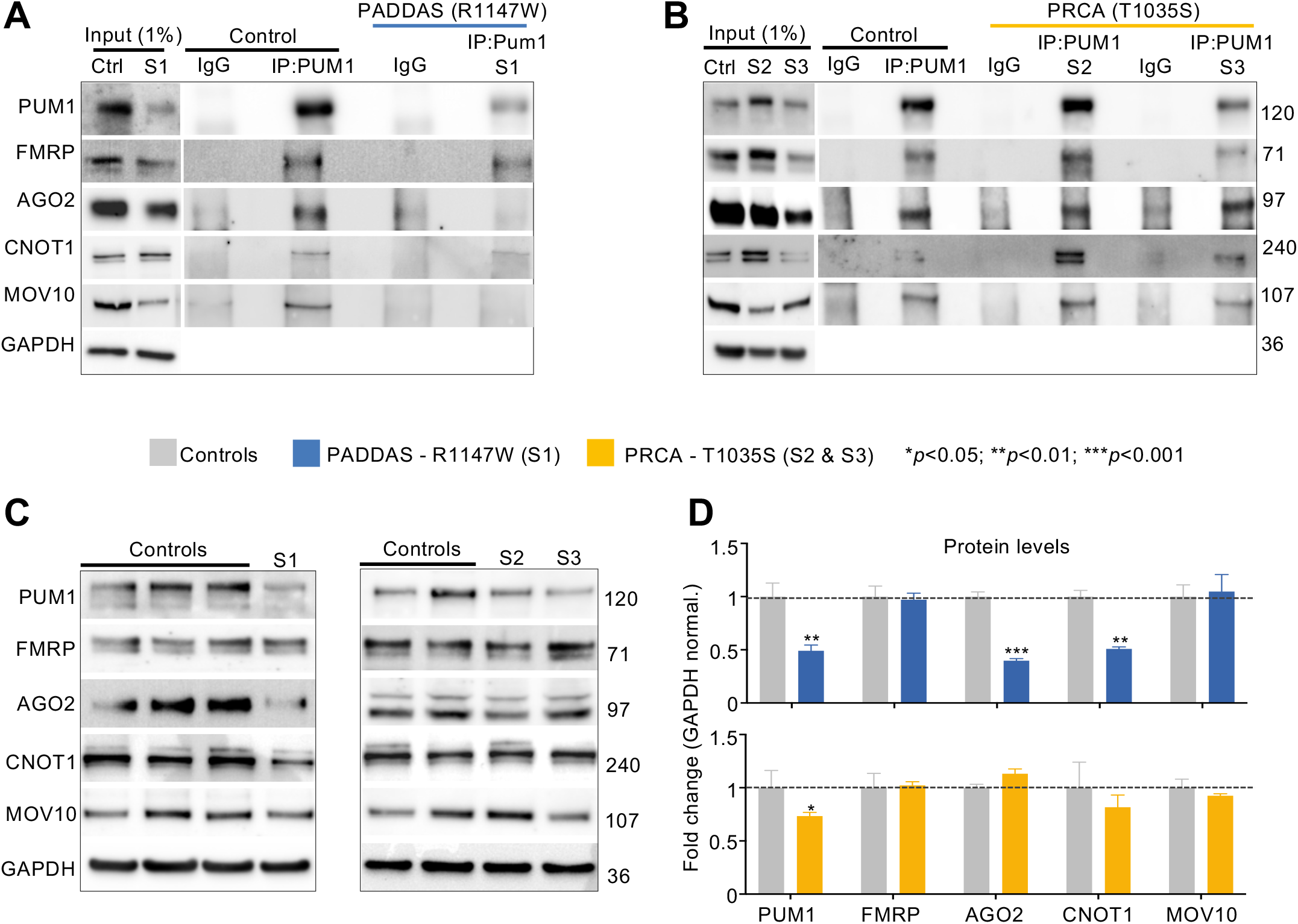
The R1147W mutation, but not T1035S, destabilizes PUM1 interactors. (**A-B**) IP against PUM1 from Subject 1 (S1, R1147W) (**A**) and Subject 2 (S2, T1035S) (**B**) compared to their respective age-, sex-, and cell type-matched healthy controls confirm the interactions between PUM1 (used here as a positive control), and PUM2, FMRP, AGO2, CNOT1, and MOV10. Input (1%) was used as a loading control and IP against IgG was used as a negative control. (**C**) Representative western blots (*left panels*) and relative quantification (*bar graphs to the right*) of protein levels for PUM1, PUM2, FMRP, AGO2, CNOT1, and MOV10 in PADDAS patient-derived and PRCA patient-derived cells compared to their respective controls. Data were normalized to Gapdh protein levels. From **A** to **C,** all the experiments were performed at least three times. Data represent mean ± SEM. P values were calculated by two-tailed Student’s t test. \**p* < 0.05, \*\**p* < 0.01, \*\*\**p* < 0.001.

Co-IP confirmed that PUM1 associates with FMRP, AGO2, CNOT1, and MOV10 in patient cell lines (Figure 6A, B). (RBFOX3 and PUM2 were expressed at too low levels in the patient-derived cells.) The mild T1035S variant had no effect on any of these proteins (Figure 6B), but R1147W impaired PUM1 association with AGO2, MOV10, and CNOT1 (Figure 6A).

In order to compare the effects of the mutants in the same cell type, we turned to HEK293T cells. GST-AGO2 associated with Myc-PUM1-R1147W 72% less than it did with Myc-PUM1-WT (Figure S13A), in alignment with our observations in the PADDAS cell lines (Figure 6A, B). CNOT1 interactions were diminished by ∼35% (Figure S13B). We observed no decrease in PUM1 association with FMRP (Figure S13C), again in accord with our findings in patient-derived cells.

We next asked whether the R1147W mutation might be impaired in binding to WT PUM1. Myc-PUM1-R1147W pulled down 51% of total GST-PUM1-WT, but the interaction between Myc-PUM1-T1035S and GST-PUM1-WT remained unchanged (Figure S13D). The same results were observed after RNase treatment (Figure S13E). Moreover, the combination of lower protein levels and marked protein instability explains why the R1147W human phenotype is closer to that of the *Pum1* null mice than to the heterozygous mice (Gennarino *et al*., 2018).

To confirm that R1147W destabilizes these PUM1 interactors, we measured these RBPs in the patient-derived cell lines. The proteins that lose their association with the R1147W variant also are reduced in their expression (Figure 6C, D). Notably, MOV10’s association with R1147W was greatly reduced (Figure 6A) even though its protein levels were unchanged (Figure 6C, D). AGO2 and CNOT1 levels were unchanged in the T1035S cell line but were ∼50% lower in the R1147W cell line (Figure 6C, D). The mRNA levels of *PUM1*, *AGO2*, *CNOT1*, and *MOV10* did not change (Figure S13F), confirming that the reductions in their respective protein levels were due to the loss of interaction with PUM1-R1147W. Collectively, these data suggest that the R1147W variant exerts a dominant-negative effect on PUM1-RBP interactors by destabilizing them.

### Shared targets are upregulated in PADDAS but not PRCA

We next tested the effects of the T1035S and R1147W mutations on both shared targets and validated PUM1-specific targets (Chen *et al*, 2012; Gennarino *et al*., 2015; Zhang *et al*., 2017) that are not in the HITS-CLIP data for the other RBPs but are expressed in both fibroblasts and lymphoblasts. PUM1-specific mRNA were dysregulated to very similar extents in PRCA and PADDAS patient cells, with only a few targets being upregulated by as much as 20% in the adult-onset disease (Figure S14A).

Of the 175 targets shared between PUM1, PUM2, AGO2, FMRP, and RBFOX3 (**Table S5**), 54 were expressed in both R1147W fibroblasts and T1035S lymphoblastoid cells. Fifty-one of those were upregulated in the former but not the latter (Figure 7A), by an average of two-fold (ranging from a low of 121% for *IDS* to 347% for *TLK1*). There was little or no change in most of these targets in T1035S cells, though levels of *CALM1*, *ATP2A2*, *CREB1*, and *GNAQ* fell by ∼40%, and *CALM2*, *TAOK1*, and *UBE2A* by ∼20% (Figure 7A).

**Figure 7.**
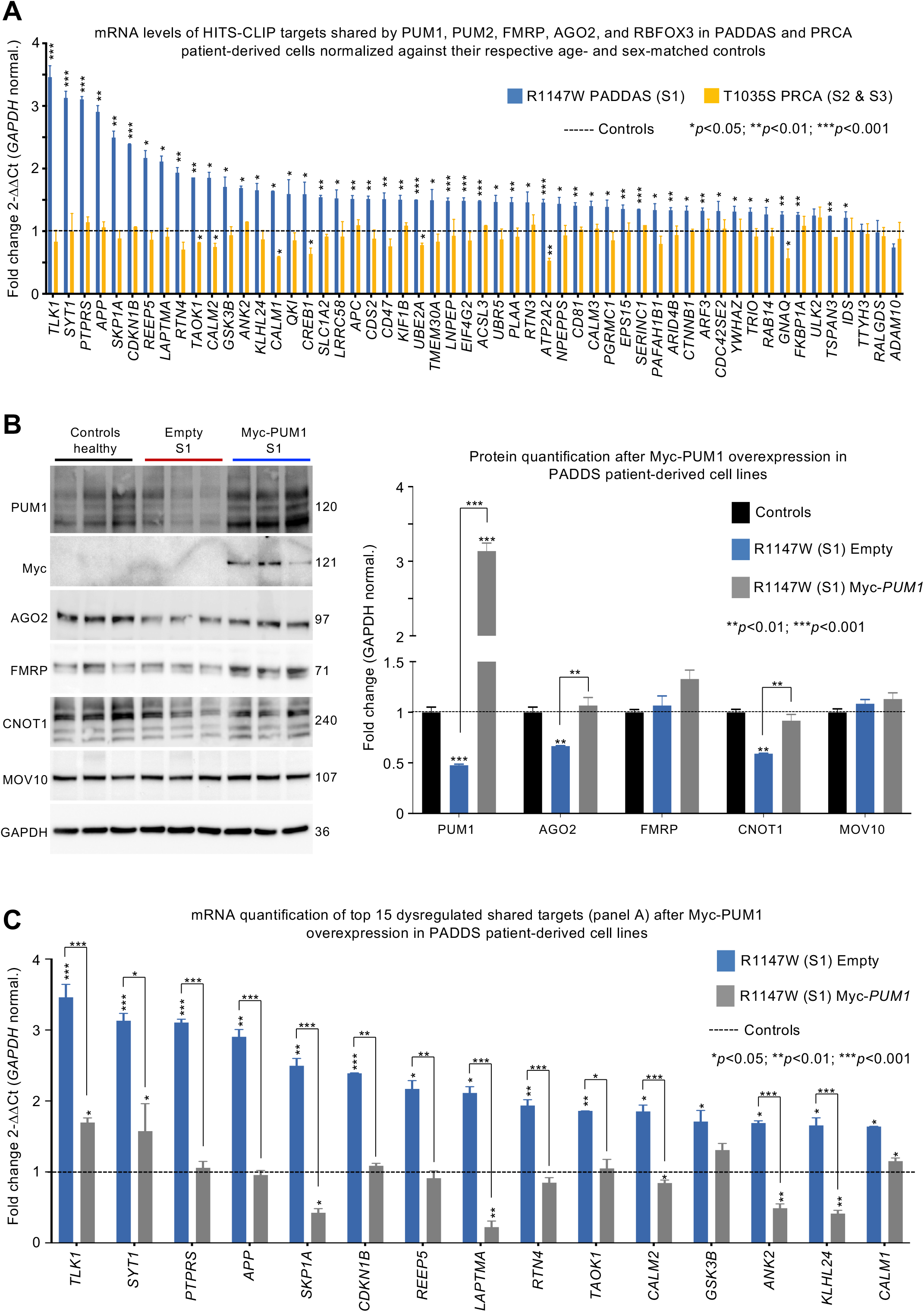
Shared targets are upregulated only in PADDAS, not in PRCA. (**A**) mRNA level quantification of PUM1 neuronal targets in common with FMRP, PUM2, AGO2, and RBFOX3 (Figure 5E and Table S5) in S1 (R1147W, *blue bars*) and S2 and 3 (T1035S, *orange bars*) cell lines compared to their respective healthy controls. Only genes expressed in both R1147W and T1035S cell lines are represented here, for a total of 54 genes. (**B**) Representative western blots (*right panel*) and relative quantifications (*left panel*) of PUM1 and its interactors (AGO2, CNOT1, FMRP, and MOV10) in R1147W patient-derived cell lines after Myc-PUM1-WT expression. (**C**) mRNA quantification of the top 15 shared target genes from panel **A** in R1147W patient-derived cell lines after Myc-PUM1-WT expression. All data were normalized to GAPDH mRNA or protein levels and experiments were performed at least three times. Data represent mean ± SEM. P values were calculated by two-tailed Student’s t test. \**p* < 0.05, \*\**p* < 0.01, \*\*\**p* < 0.001. The full list of shared targets expressed in fibroblast and lymphoblast cell lines is reported in Table S5.

To determine whether restoring PUM1 levels would normalize expression of these shared targets, we transfected Myc-PUM1-WT in R1147W cells, using transfection with an empty vector as a negative control (Figure 7B, Figure S14B). WT PUM1 rescued AGO2 and CNOT1 protein levels compared to age- and sex-matched healthy controls and this reduced the levels of the top 15 upregulated shared targets (Figure 7B, C). These data confirm that the effects of the R1147W mutation, which does not impair PUM1 binding to mRNA (Gennarino *et al*., 2018), result from loss of interactions with RBPs that repress the same mRNA targets. These results also support the hypothesis that the symptoms observed in adult-onset SCA47 are attributable to the dysregulation of PUM1-specific target genes, while infantile-onset SCA47 involves both protein partner destabilization and dysregulation of the partner proteins’ targets.

## Discussion

Since our initial study describing PUM1-related phenotypes (Gennarino *et al*., 2018), we and others have identified additional SCA47 patients (Bonnemason-Carrere *et al*, 2019; Gennarino *et al*., 2018; Imaizumi *et al*, 2019; Lai *et al*, 2019). In our cohort, heterozygous deletion and the R1147W mutation account for the majority of infantile-onset patients, and T1035S for the majority of adult-onset cases. The question that drove the present study is: why should the additional 25% drop in PUM1 levels from PRCA to PADDAS produce such different phenotypes, especially when R1147W is not impaired in binding to mRNA? Our data support the hypothesis that the difference is not due to a linear increase in the de-repression of mRNA targets but is rather a function of an additional mechanism coming into play: the destabilization of numerous interactors and the de-repression of their downstream targets. This conclusion relies on five lines of evidence. First, loss of Pum1 in heterozygous and knockout mice changes the levels of associated proteins, with unexpected differences emerging between brain regions and between male and female mice. The odds of consistently identifying specific proteins in different brain regions and sexes as false positives, across as many mice as these experiments required, are extremely low. Second, we observed diminished function of the RBP interactors in the absence of Pum1, insofar as their targets are dysregulated in Pum1-KO mice; moreover, the dysregulation of miRNA showed opposite patterns in male and female cerebella that correlated with the sex-specific patterns of Ago2 expression. Third, the levels of these proteins were reduced 40-70% in PADDAS patient cell lines, despite unaltered mRNA levels, but not in PRCA patient cells; we also found that 55 shared targets expressed in both lymphoblasts and fibroblasts were derepressed in infantile-onset SCA47, but not adult-onset SCA47, cells. Fourth, our *in vitro* studies showed that AGO2, CNOT1, and WT PUM1 lose their interaction with PUM1-R1147W. Fifth, re-expression of PUM1 in PADDAS cell lines restored the abundance of its interactors and suppression of downstream targets. These results underscore the importance of examining RBP interactions *in vivo*, in specific contexts (different sex or brain regions), with and without RNase treatment.

There are other dosage-sensitive proteins that produce different phenotypes depending on their expression level (Tan & Zoghbi, 2019), and our results raise the possibility that interacting complexes may be disrupted once expression falls below a certain threshold, which for PUM1 appears to be around 50%. It is worth noting that a recent study found that, below a threshold of ∼70% of normal levels of FMRP, there were steep decreases in IQ for each further decrement in FMRP levels, even as small as 5-10% (Kim *et al*, 2019). The amount of loss that can be sustained for a given protein would likely depend on its usual abundance, and it is possible that for some proteins, the difference in phenotype between greater and lesser abundance reflects a linear progression from mild to severe. For example, in proteopathies such as Alzheimer’s or Parkinson’s disease, genetic duplications of *APP* or *SNCA* cause an earlier onset of what is recognizably the same disease (Chartier-Harlin *et al*, 2004; Rovelet-Lecrux *et al*, 2006). Similarly, a mutation in *MECP2* that reduces its protein levels by only 16% is still sufficient to cause a mild form of Rett syndrome (Takeguchi *et al*, 2020).

In contrast, there are diseases in which the phenotypes do not simply range from mild to severe versions of the same symptoms, but seem to take on a different character. In the polyglutaminopathies, the disease-causing protein bears an abnormally expanded CAG tract that tends to expand upon intergenerational transmission. Although the range of normal and pathogenic repeat tract lengths differs from one polyglutamine disease to another, larger expansions are more unstable, cause earlier onset, and affect far more tissues than smaller expansions (Orr & Zoghbi, 2007). For example, adult-onset SCA7 presents as ataxia, but infantile SCA7 affects the entire nervous system, the heart, and the kidneys, and leads to death by two years of age (Bah *et al*, 2020). Another example is Huntington’s disease (HD), where the juvenile form frequently lacks the classic chorea yet produces seizures, which are not a feature of the adult-onset disease; brain morphometry is also quite different in adult- and juvenile-onset cases (Tereshchenko *et al*, 2019). In this family of diseases, therefore, the mechanism is the same (repeat expansion), but different tissues have different thresholds for the CAG repeats. Moreover, the brain regions most vulnerable to HD show dramatic levels of somatic instability that correlate better with clinical outcomes than the germline polyglutamine expansion (Genetic Modifiers of Huntington’s Disease, 2015; Kacher *et al*, 2021).

In the case of PUM1-related disease, it seems that an additional mechanism comes into play for the more severe phenotypes, beyond upregulation of mRNA targets. Interestingly, FMRP, which harbors a dynamic CGG repeat, is also associated with two very different phenotypes, through two different mechanisms: very large expansions silence the gene and produce Fragile X syndrome, whereas premutations cause the adult-onset Fragile-X-associated tremor/ataxia syndrome through RNA toxicity (FXTAS) (Salcedo-Arellano *et al*, 2020). Interestingly, the clinical presentation of FXTAS differs by sex.

Our data suggest that the three mechanisms of repression that have been proposed for PUM1— collaborating with the miRNA machinery (Friend *et al*., 2012; Kedde *et al*., 2010; Miles *et al*., 2012), recruiting the CCR4-NOT deadenylase complex to trigger degradation (Temme *et al*., 2014; Van Etten *et al*., 2012; Weidmann *et al*., 2014), and antagonizing poly(A)-binding proteins to repress translation (Goldstrohm *et al*., 2018)—might be coordinated in neurons, insofar as PUM1, PUM2, FMRP, AGO2, MOV10, CNOT1 and RBFOX3 (and related proteins in specific brain regions) either interact or are so close to each other within the ribonucleosome that the loss of Pum1 or RNA can change the composition of the complexes that are identified by co-IP, in ways that are specific to brain region and sex. In this context it is worth noting that a very recent study found alternative splicing is altered in hippocampal slices from Fmrp-deficient mice; this observation was attributed to changes in H3K36me3 levels (Shah *et al*, 2020), but our data suggest that FMRP has a closer relationship with the RBFOX protein family and alternative splicing machinery than previously imagined. Indeed, recent work has provided tantalizing glimpses of close interactions among various kinds of RNA metabolism. For example, members of the RBFOX family of proteins may, depending on their interactors (and perhaps cell type, sex, age, and species), be involved in microRNA processing in the nucleus and translation in the cytoplasm (Conboy, 2017). The FMRP/MOV10 complex appears to be involved in regulating translation through miRNA, with evidence that this role may change according to cell type (Kenny *et al*., 2020). Another study used quantitative mass spectroscopy to examine how Fmrp expression levels change with age in the wild-type rat dentate gyrus and found differences in the levels of myriad proteins; among the 153 proteins with the most significant changes in levels were Pum1, Pum2 and Papbc1 (Smidak *et al*, 2017). The region-, sex-, and age-specificity of certain interactions indicates that unraveling RBP interactomes *in vivo* will require considerable finesse. But creating such interactomes should lead to a more complex yet realistic picture of RBP roles in neuronal function and in neurological disease.

## Acknowledgments

We thank the patients and their families for participating in our study. We thank Mingyu Cao for help with statistical analyses, U.N. Wasko for starting the cloning procedure with Myc and GST tag sequences *in vitro*, Wayne Miles for sharing the RIP protocol and send us the HCT116-*PUM1^-/-^* cell lines, and V. L. Brandt for editing the manuscript and suggesting we examine the targets of PUM1 interactors. We also thank G. Struhl, V. Ambros, T. Duchaine, G. Karsenty, H. Zoghbi, M. Jovanovic, Y. Giardina, and members of the Gennarino laboratory for helpful discussions and support. This work was supported by the National Institute of Neurological Disorders and Stroke (NINDS; R01NS109858 to V.A.G.); the National Ataxia Foundation/Young Investigator Research Grant (V.A.G.); the National Institute of General Medical Sciences (NIGMS; R35GM128855 to N.G.B.); the National Human Genome Research Institute (NHGRI; R01HG012216 to M.J.); the National Institute on Aging (NIA; R01AG071869 to M.J.); the National Institute of General Medicine (NIGM; R35GM128802 to M.J.); the Brain & Behavior Research Foundation Young Investigator Award (V.A.G.); the Paul A. Marks Scholar Program, Columbia University Vagelos College of Physicians and Surgeons (V.A.G.); the TIGER Award, Taub Institute, Columbia University Irving Medical Center (V.A.G.); the Columbia Stem Cell Initiative (CSCI), Columbia University Irving Medical Center (V.A.G. and N.dP.). The authors declare no competing interests.

## Author contributions

S.B. designed and performed molecular experiments, analyzed and interpreted the data, and drafted the manuscript. N.D.P. performed the qPCR experiments, *in vitro* immunoprecipitation assays, analyzed the data, and edited the manuscript. A.C. performed all the brain region dissections in mice, performed the qPCR experiments and maintained the *Pum1* mutant mouse colony. M.C. performed the cloning experiments. P.P and R.K.S. performed mass spectrometry and helped analyze the data. C.J.T. and N.G.B. developed the recombinant proteins. M.J and E.D.M performed and analyzed the proteomics studies. V.A.G. conceived the study, analyzed and interpreted the data, performed molecular experiments, and wrote the manuscript.

## Declaration of interests

The authors declare not competing interests

## Supplemental figures and legends

**Figure S1.**
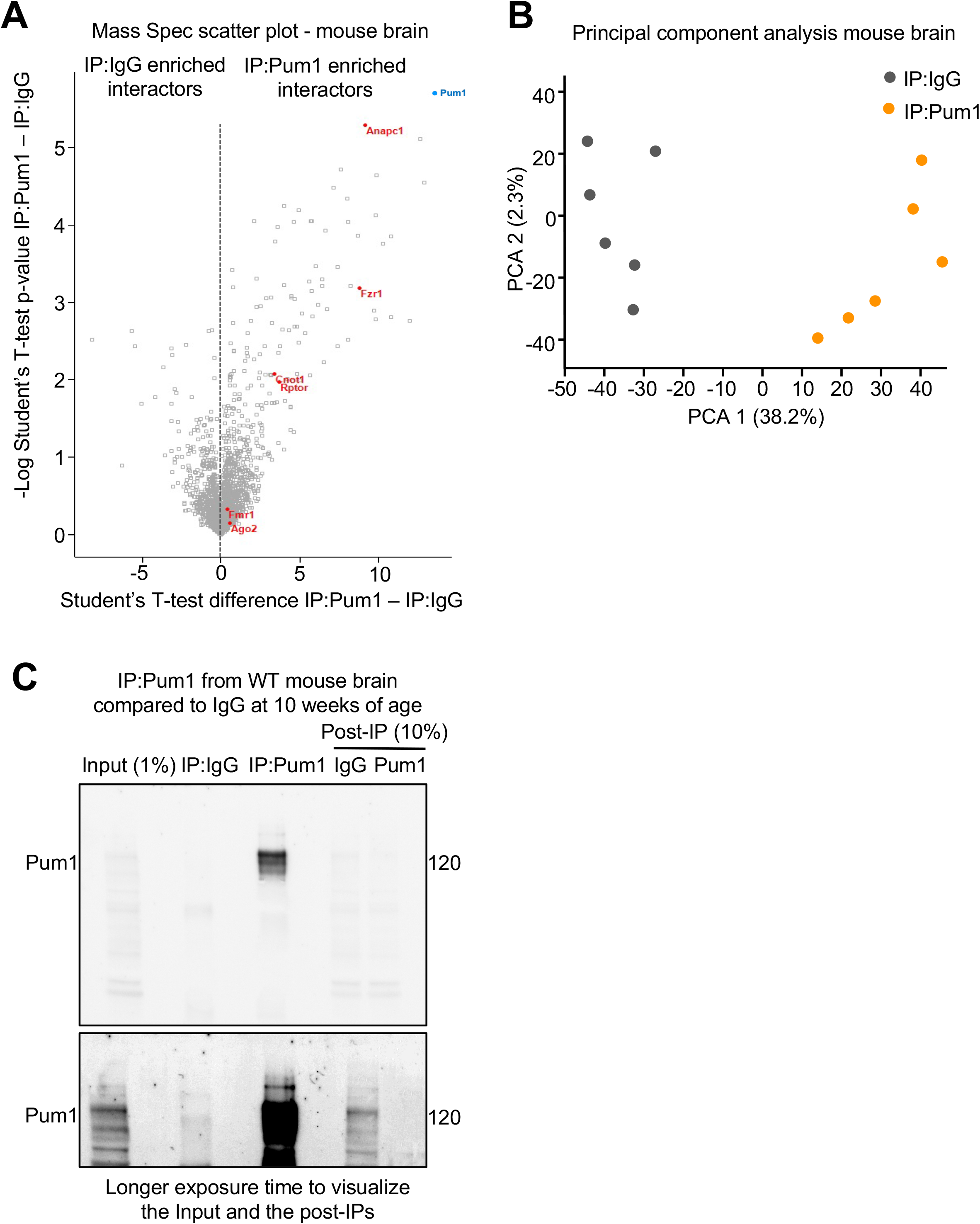
Pumilio1 antibody efficiency for IP LC-MS/MS. (**A**) Volcano plot analysis showing all the proteins pulled down by IP against IgG and Pum1 from mouse brain. (**B**) Principal component analysis (PCA) of IP-Pum1 followed by LC-MS/MS in WT adult mouse brains compared with IP against IgG; each dot represents one sample for a total of 12 samples processed by LC-MS/MS. (**C**) Pre-IP, IP, and post-IP against Pum1 and IgG from wild-type (WT) mouse brain. Even at very long exposure, the post-IP (10%) Pum1 lane has no residual band at 120 kDa even though 10 times more protein is loaded than Input (1%). This demonstrates the specificify of the Pum1 antibody, which makes it suitable for IP LC-MS/MS. The numbers on the right show molecular weight in kilodaltons (kDa).

**Figure S2.**
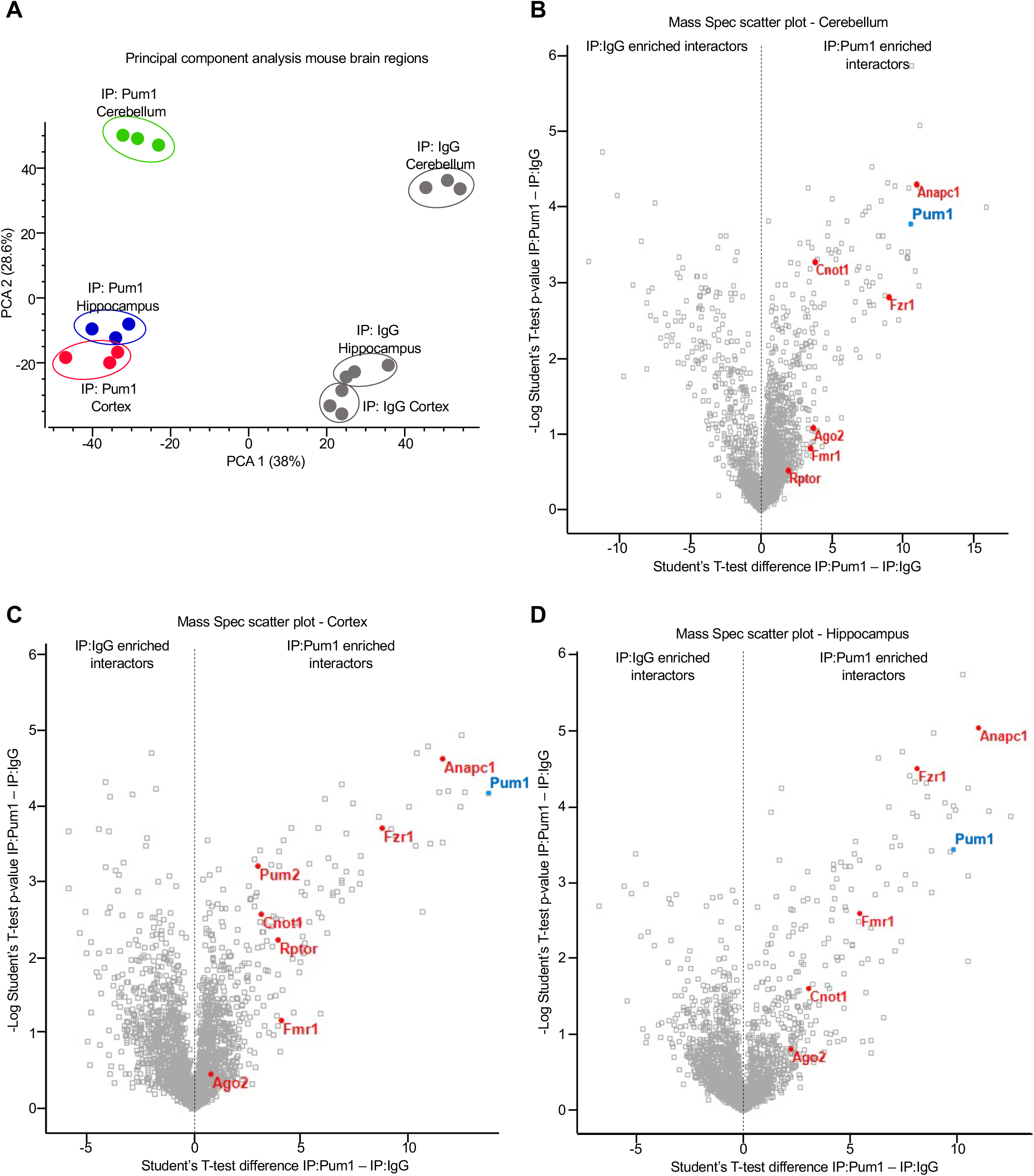
Volcano plot and PCA analyses of IP:Pum1 followed by LC-MS/MS in cerebellum, hippocampus, and cortex. **(A)** Principal component analysis (PCA) of IP-Pum1 followed by LC-MS/MS in cortex, hippocampus, and cerebellum from WT mice. (**B**-**D**) Volcano plots show all the proteins pulled down by IP against IgG and Pum1 from (**B**) cerebellum, (**C**) cortex, and (**D**) hippocampus at 10 weeks of age. IP against IgG was used as a negative control. Each dot represents a total of 3 samples processed by MS for each brain region. All putative Pum1 interactors are listed in **Table S1**.

**Figure S3.**
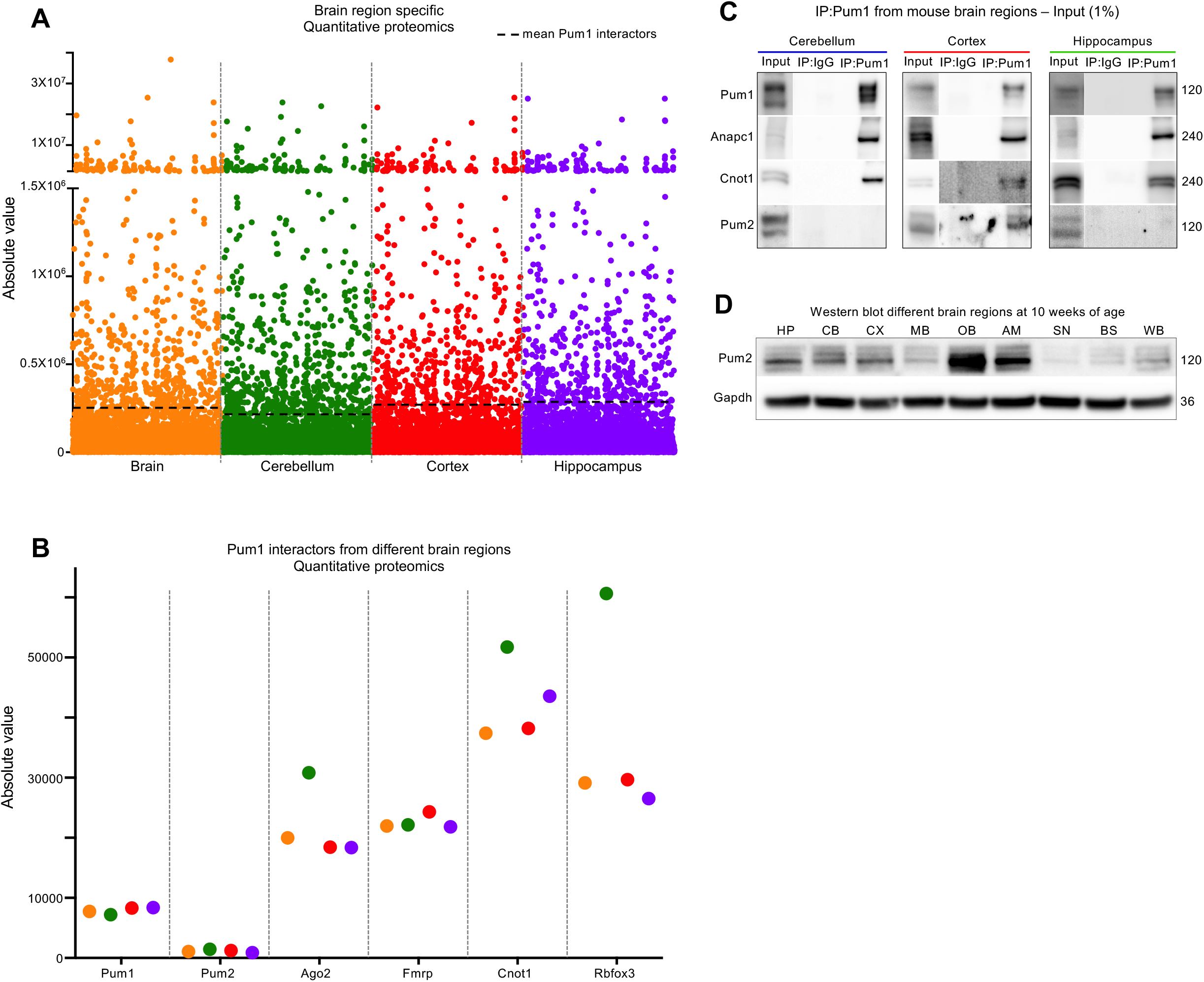
Many Pum1 interactors are specific to certain brain regions and not determined by expression level. (**A**) Proteomic analysis in whole brain, cerebellum, hippocampus, and cortex at 10 weeks of age shows that Pum1 interactors are not the most highly expressed proteins. Dotted line represents the mean expression of Pum1 interactors from each brain region. Proteomics was performed in duplicate (one male and one female) for each brain region. (**B**) Absolute expression value of the validated Pum1 interactors. Pum2 is expressed a low levels in all three brain regions but was still the strongest Pum1 interactor in the cortex, suggesting a specific interaction. (**C**) Immunoblot for Pum1 (positive control), Anapc1, Cnot1 and Pum2. (**D**) Western blot analysis at 10 weeks of age to evaluate Pum2 expression levels in eight different brain regions as well as whole brain. Pum2 is highly expressed in olfactory bulbs and amygdala, and expressed at similar levels in hippocampus, cerebellum, and cortex. All experiments performed in triplicate. Cerebellar and cortical tissues: n=8 wild-type mice (4 male and 4 female), for a total of 24 mice. Hippocampus: n=10 wild-type mice (5 female and 5 male), for a total of 30 mice. All mice were 10 weeks of age. IP against IgG was used as a negative control. Molecular protein weights are expressed in kilodaltons (kDa). HP: hippocampus; CB: cerebellum; CX: cortex; MB: midbrain; OB: olfactory bulbs; AM: amygdala; SN: substantia nigra pars compacta; BS: brain stem; WB: whole brain. All the experiments were repeated at least three times.

**Figure S4.**
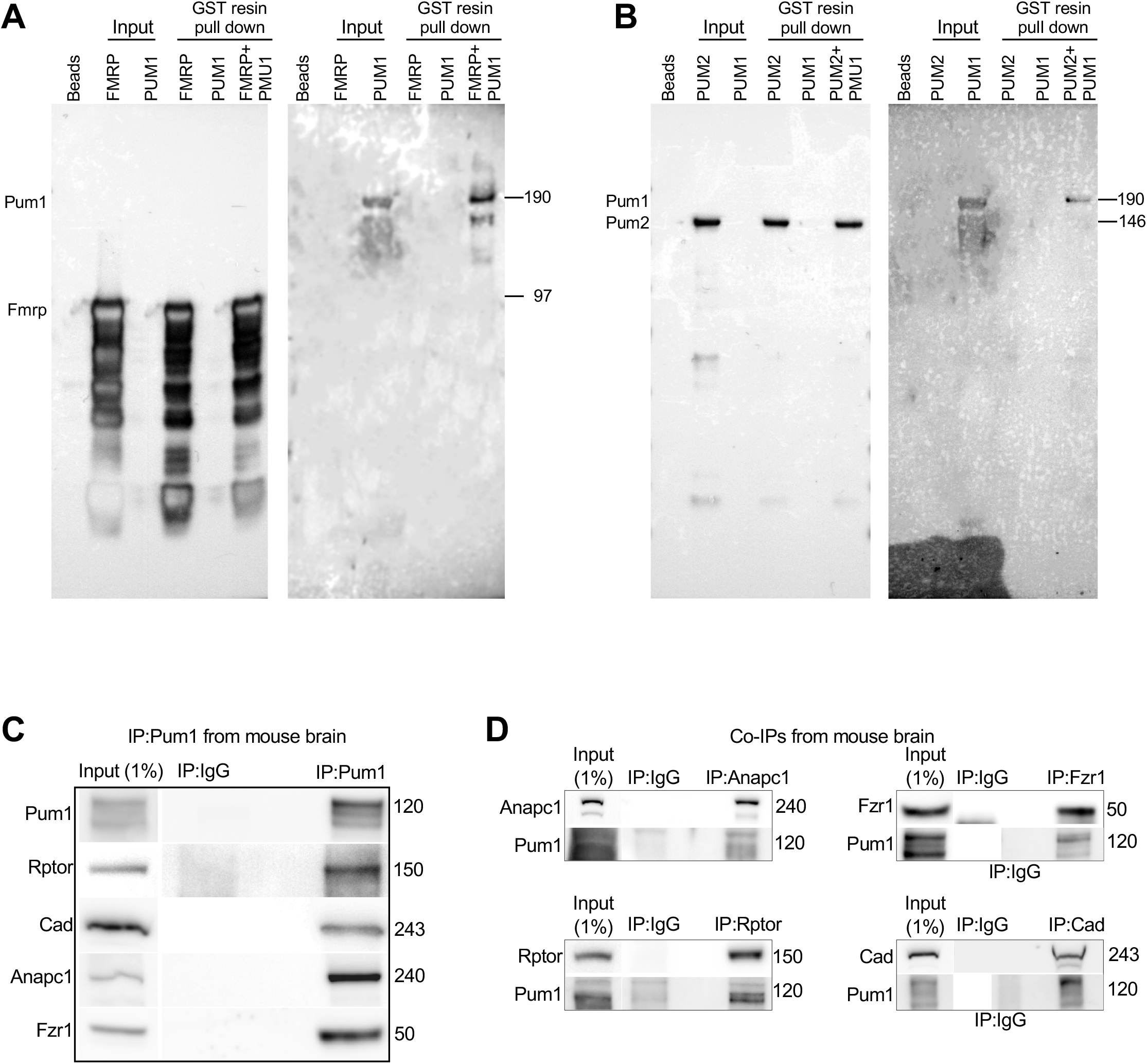
Fmrp and Pum2 IP recombinant proteins and validation of APC/C and mTOR proteins. (**A-B**) Representative western blot of PUM1 pulled down by recombinant (**A**) GST-FMRP or (**B**) GST-PUM2. (**C**) Representative western blot of Rptor, Cad, Anapc1 and Fzr1 proteins pulled down by IP against Pum1 (used here as positive control) from WT mice. (**D**) Representative western blot of the reciprocal co-IPs against Anapc1, Fzr1, Rptor, and Cad. Each co-IP was immunoblotted against Pum1 and the pulled down protein used here as a reference protein to confirm the respective protein-protein interaction. For **C** and **D**, IP against IgG was used as a negative control, and Input (1% from the initial protein lysate) as a loading control. Molecular weights are expressed in kilodaltons (kDa) to the right. All the experiments were repeated at least four times. All mice were sacrificed at 10 weeks of age with an equal number or male and female.

**Figure S5.**
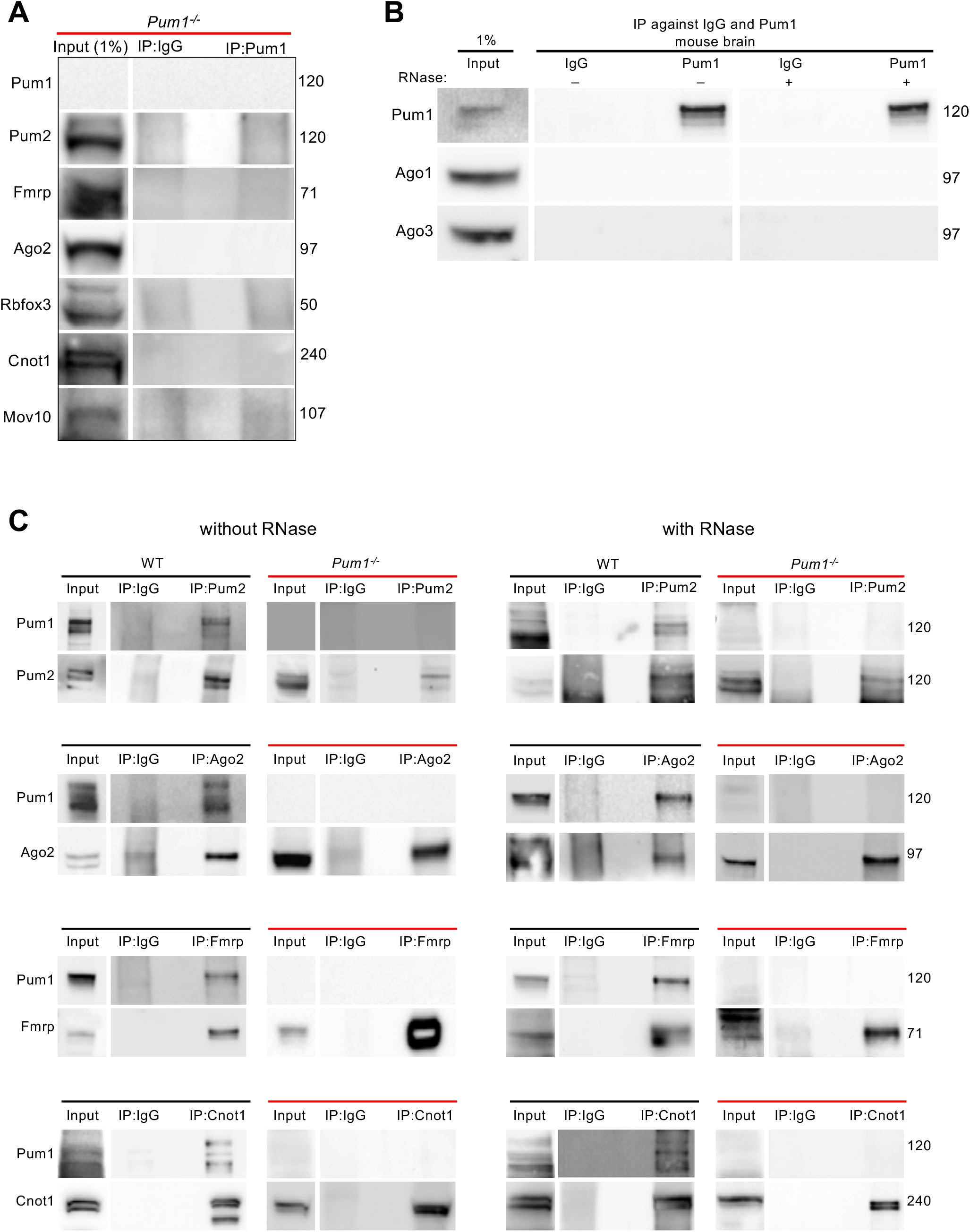
IP and co-IP against Pum1 and the interactors from whole brain in *Pum1^-/-^*and WT mice, with and without RNase. (**A**) IP against Pum1 in *Pum1^-/-^* mouse demonstrates the complete absence of Pum1 and thus the specificity of the anti-Pum1 antibody. IP against IgG was used as a negative control, and Input (1% from the initial protein lysate) as a loading control. (**B**) IP against Pum1 (with or without RNase treatment) shows no interaction with Ago1 or Ago3 in the mouse brain. (**C**) Representative western blots of Pum1 pulled down by Pum2, Fmrp, Ago2, Cnot1 from WT and *Pum1^-/-^* mouse brains, with and without RNase. Molecular weights at the right are in kilodaltons (kDa). All the experiments were conducted with one female and one male mouse brain at 10 weeks of age, in triplicate, for a total of 6 mice. IP against IgG was used as a negative control, and Input (1% from the initial protein lysate) as a loading control.

**Figure S6.**
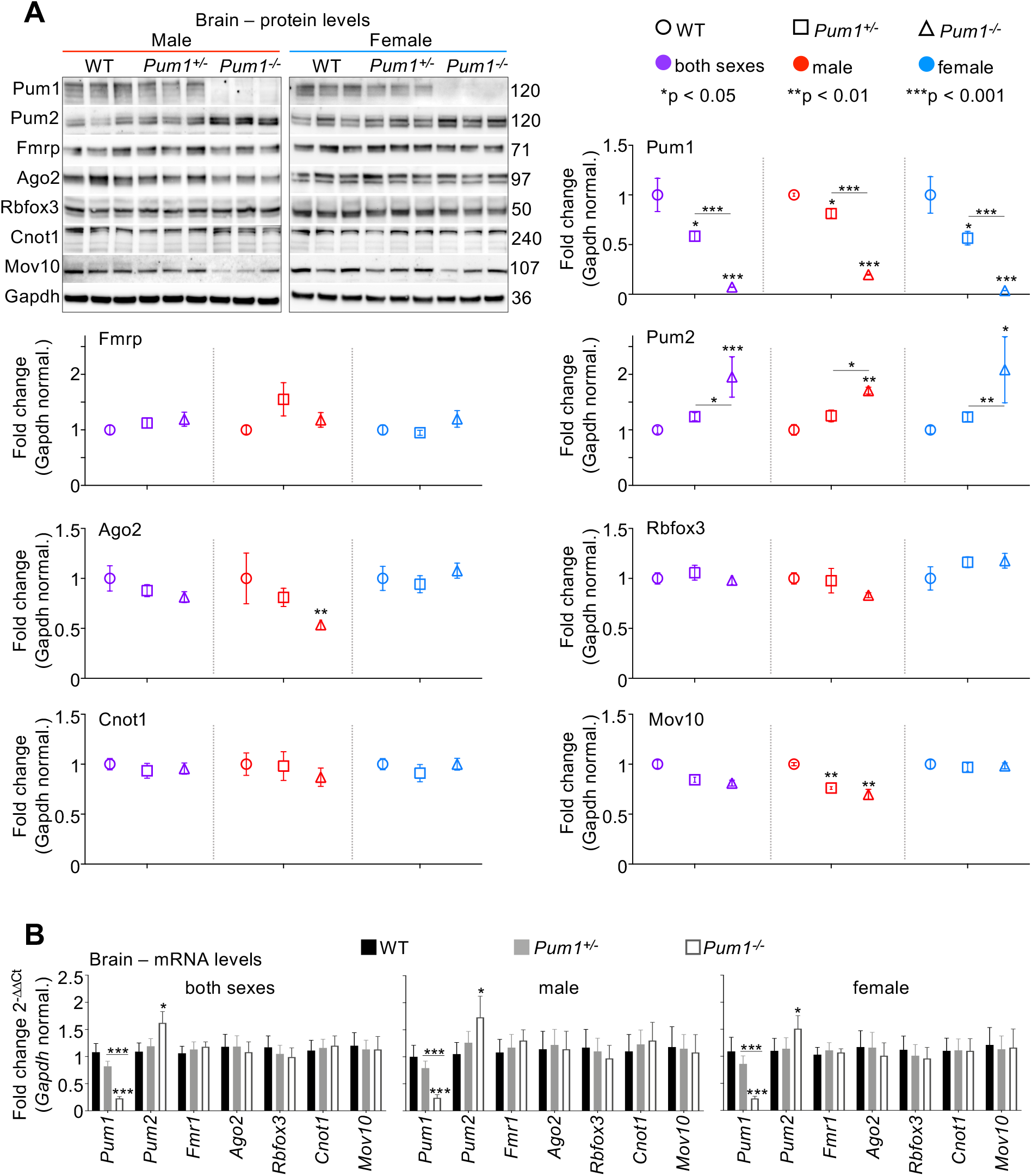
Protein and mRNA quantification from WT, *Pum1^+/-^* and *Pum1^-/-^* mouse brains. (**A**) Representative western blot with relative quantifications of Pum1, Pum2, Fmrp, Ago2, Rbfox3, Cnot1, and Mov10 from whole brains of WT, *Pum1^+/-^* and *Pum1^-/-^* mice. All data were normalized to Gapdh protein levels. The numbers on the right are the respective molecular weights expressed in kilodaltons (kDa). (**B**) mRNA level quantification by qPCR of *Pum1*, *Pum2*, *Fmrp*, *Ago2*, *Rbfox3*, *Cnot1*, and *Mov10* from whole brains of WT, *Pum1^+/-^*and *Pum1^-/-^* mice. Again, all data were normalized to *Gapdh* mRNA levels. All the experiments were conducted with equal number of male (at least 6 per genotype) and female (at least 6 per genotype) mice at 10 weeks of age, for a total of at least 12 mice per genotype (data represent mean ± SEM). The *p* values were calculated by the Student’s t test. \**p* < 0.05, \*\**p* < 0.01, \*\*\**p* < 0.001.

**Figure S7.**
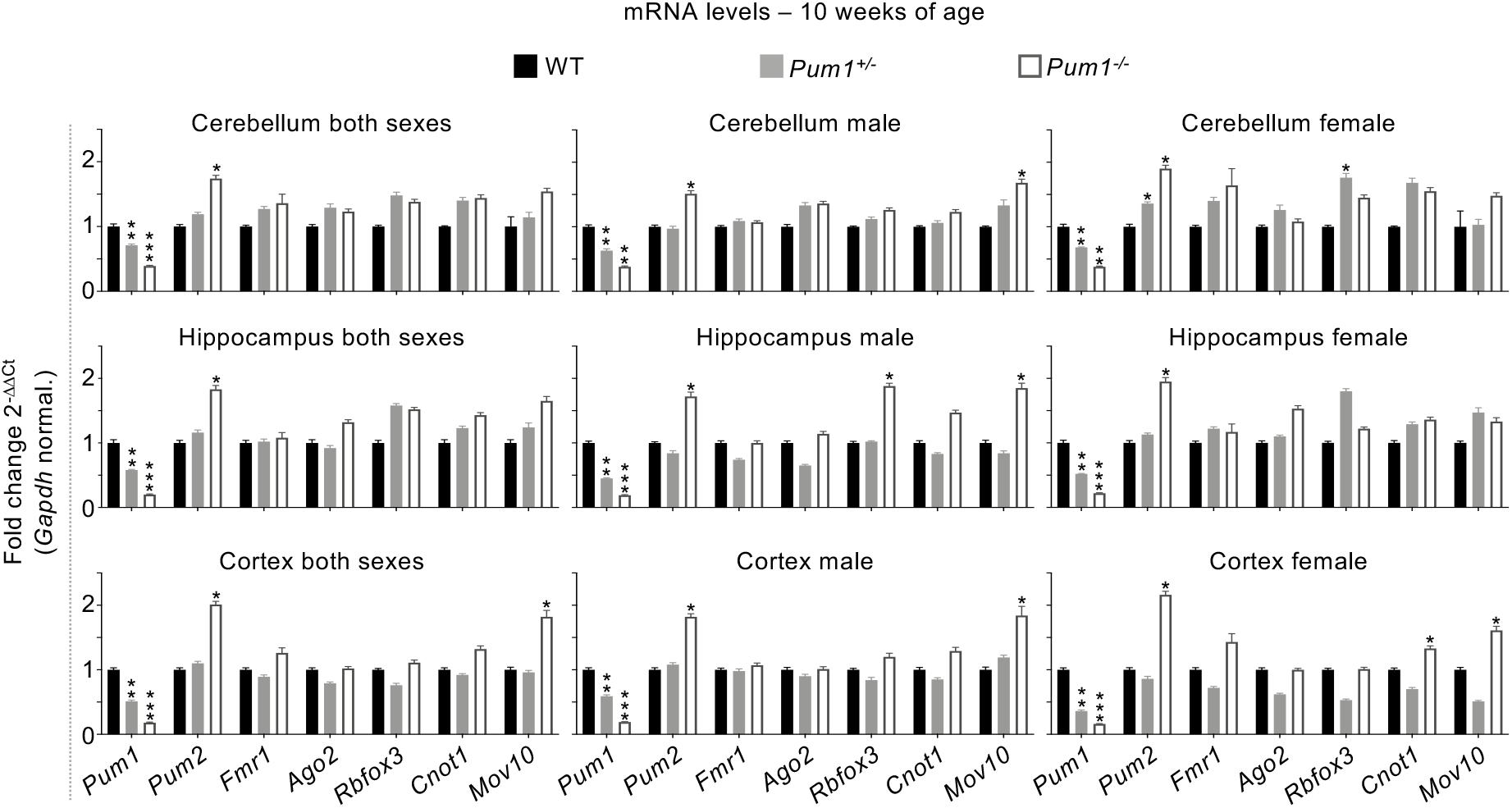
mRNA quantification of Pum1 interactors by brain region and sex in WT, *Pum1^+/-^* and *Pum1^-/-^* mice. mRNA levels in cerebellum, hippocampus, and cortex in male and female for all the validated Pum1 interactors. The same number of mice were used here as in Figure 4A-C for a total of at least 12 mice per genotype and sex at 10 weeks of age. All data were normalized to *Gapdh* mRNA levels. All the experiments were performed at least six times (data represent mean ± SEM). The *p* values were calculated by the Student’s t test. \**p* < 0.05, \*\**p* < 0.01, \*\*\**p* < 0.001.

**Figure S8.**
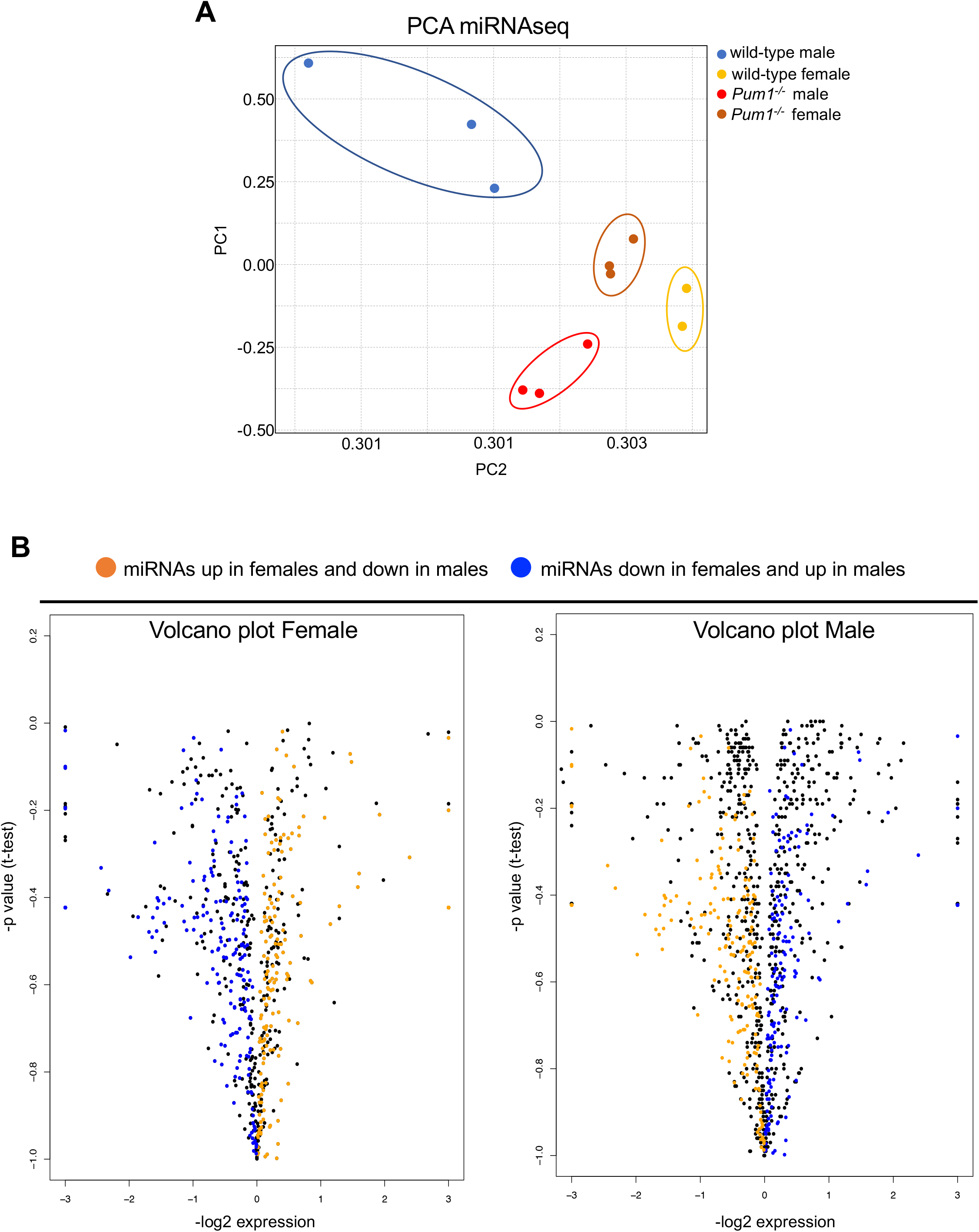
Volcano plots representing all the miRNAs sequenced by miRNAseq in male and female. (**A**) Principal component analysis (PCA) of miRNAseq in wild-type and *Pum1*^-/-^ male and female mice cerebella at 10 weeks of age. (**B**) Volcano plots show the expression profile for all the miRNAs in male and female *Pum1^-/-^* mice compared to WT at 10 weeks of age. The orange dots represent the miRNAs upregulated in female and downregulated in males; the blue dots represent the miRNAs downregulated in female and upregulated in males (see Material and Methods).

**Figure S9.**
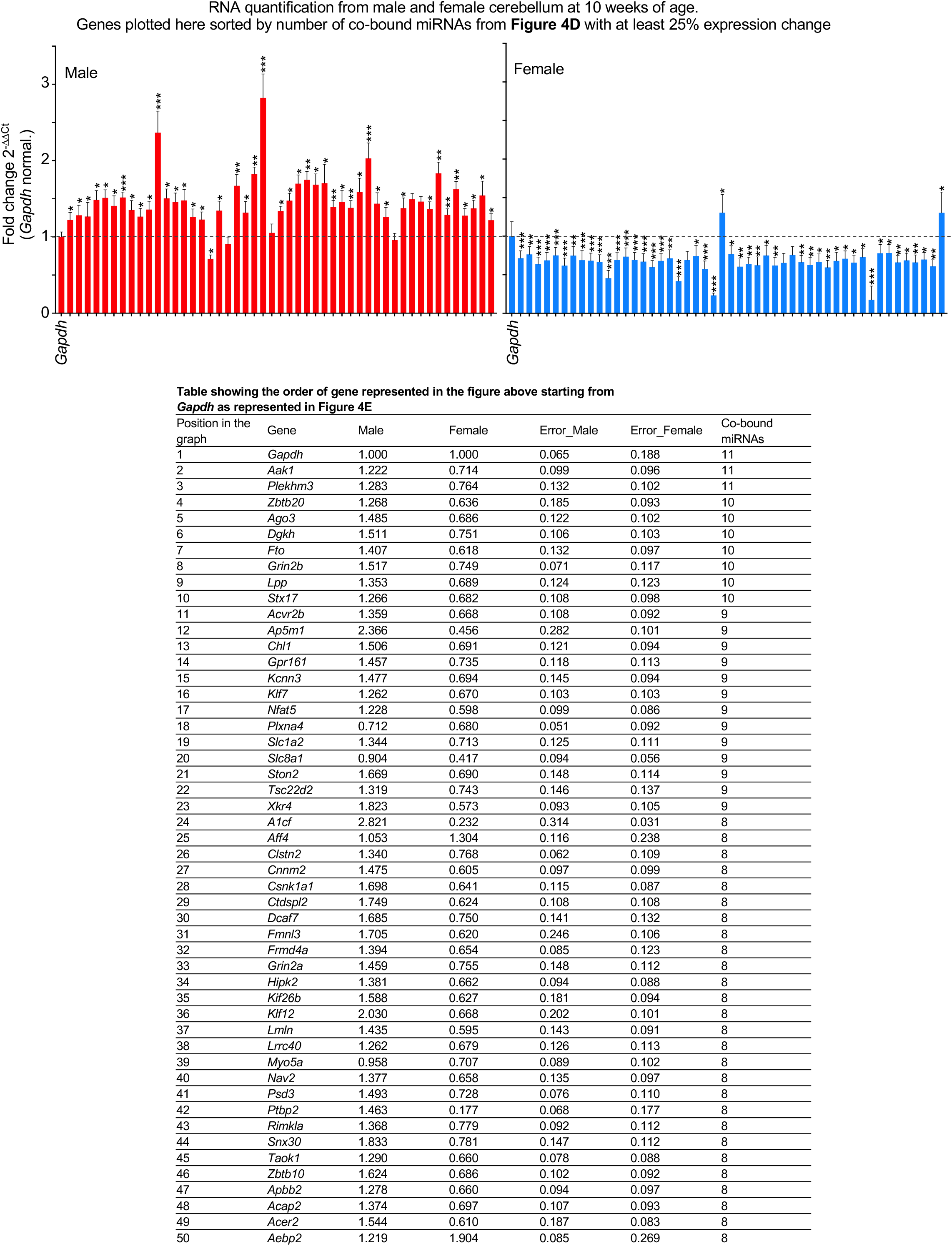
mRNA quantification of the 49 targets co-bound by at least eight dysregulated miRNAs in mouse cerebellum. qPCR in cerebellum of male (*left*, red) and female (*right*, blue) mice at 10 weeks of age for the 49 targets co-bound by at least eight dysregulated miRNAs (with minimum 25% change in expression) from Figure 4E and **Table S4**. All the experiments were performed in triplicate for both male and female (data represent mean ± SEM). The *p* values were calculated by two-tailed Student’s t test. \**p* < 0.05, \*\**p* < 0.01, \*\*\**p* < 0.001.

**Figure S10.**
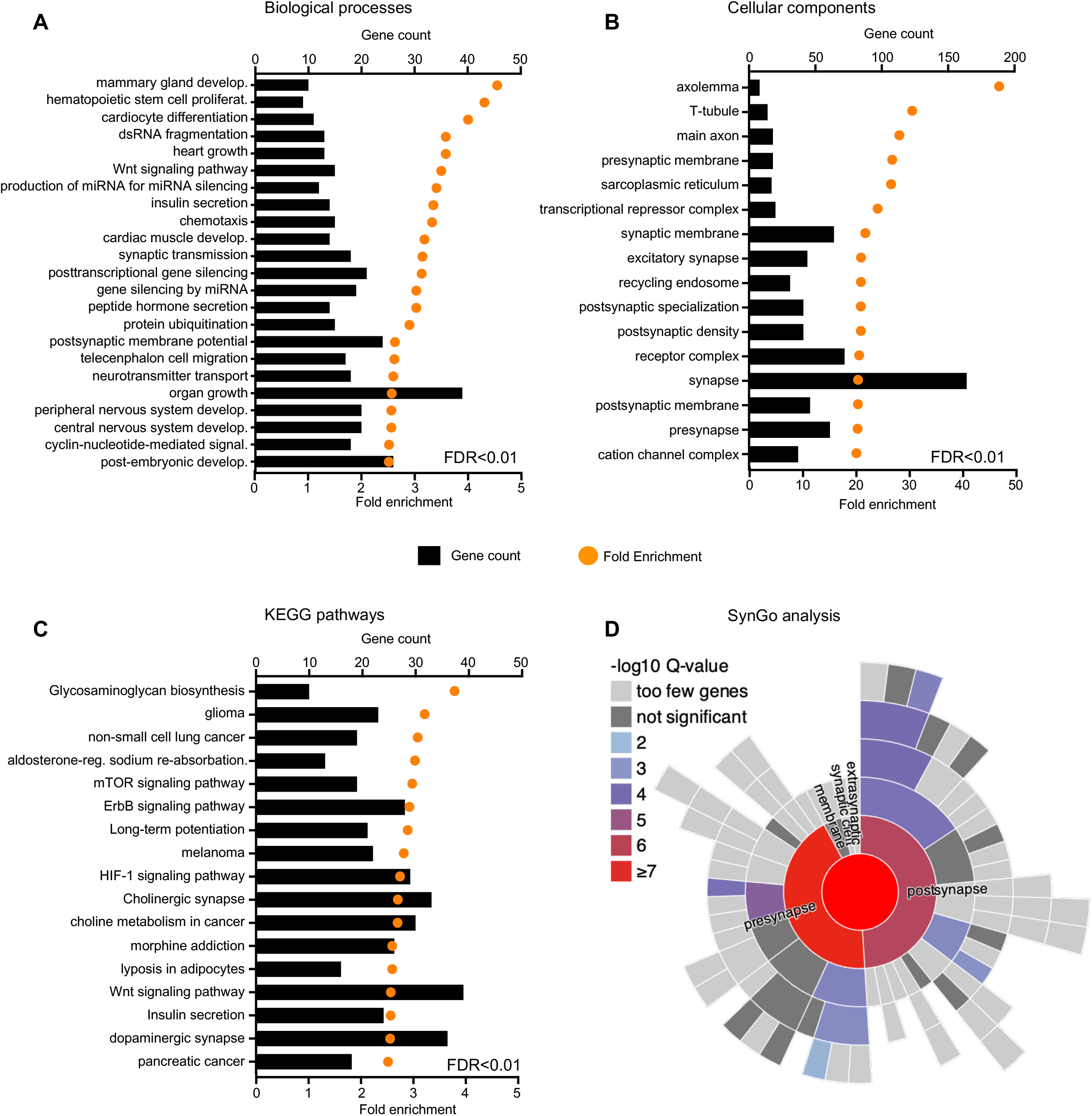
Gene ontology analysis for all targets predicted by CoMeTa and TargetScan that are co-bound by at least four miRNAs. (**A**-**C**) David Gene Ontology representing the enriched (**A**) biological processes, (**B**) cellular components, and (**C**) KEGG pathways for all the targets co-bound by at least four miRNAs. For this analysis we set FDR<0.01 and a fold-enrichment >2. (**D**) Synaptic Gene Ontology (SynGO) predicts that 117 targets are presynaptic and 124 are postsynaptic with a log10Q value≥5.

**Figure S11.**
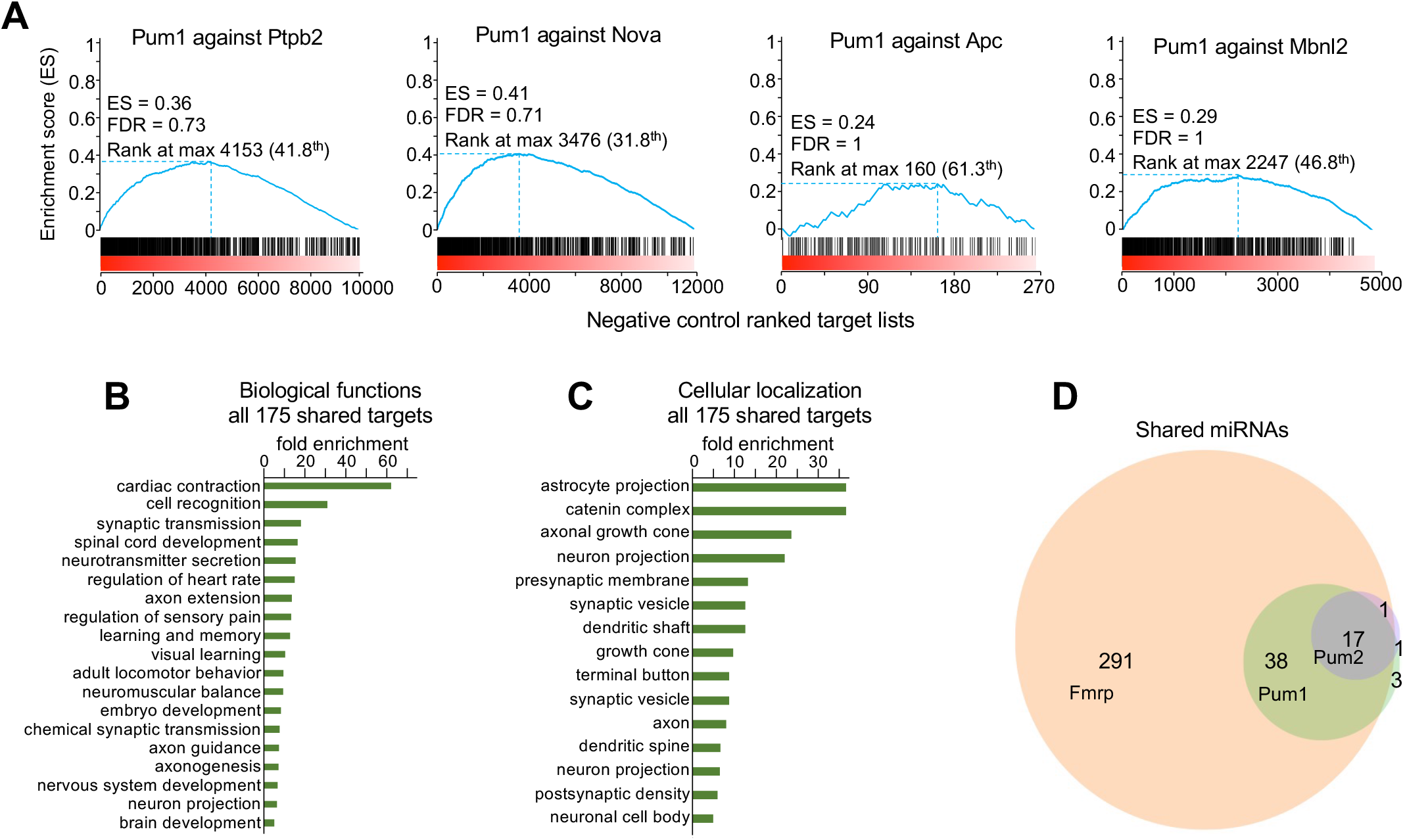
GSEA and Gene Ontology data pertaining to Figure 5. (**A**) Gene Set Enrichment Analysis (GSEA) of Pum1 HITS-CLIP data plotted against HITS-CLIP data from the negative controls (RBPs that did not show up in the Pum1 interactome: Ptpb2, Nova, Apc, and Mbnl2) reveals no significant enrichment. (**B**-**C**) Gene ontology analysis of the HITS-CLIP targets shared between Pum1, Pum2, Fmrp, Ago2, and Rbfox3 reveals enrichment for certain (**B**) biological functions and (**C**) cellular localization. Only categories with FDR<0.05 and fold enrichment > 5 were plotted in **B** and **C**. (**D**) Venn diagram of miRNAs identified by Pum1 and Pum2 shows almost 100% overlap with the miRNAs pulled down by Fmrp HITS-CLIP. For full list of shared miRNAs see **Table S6**. For all GSEA analyses the False Discovery Rate (FDR) was provided by GSEA, ***FDR < 0.01. ES=Enrichment score (blue line).

**Figure S12.**
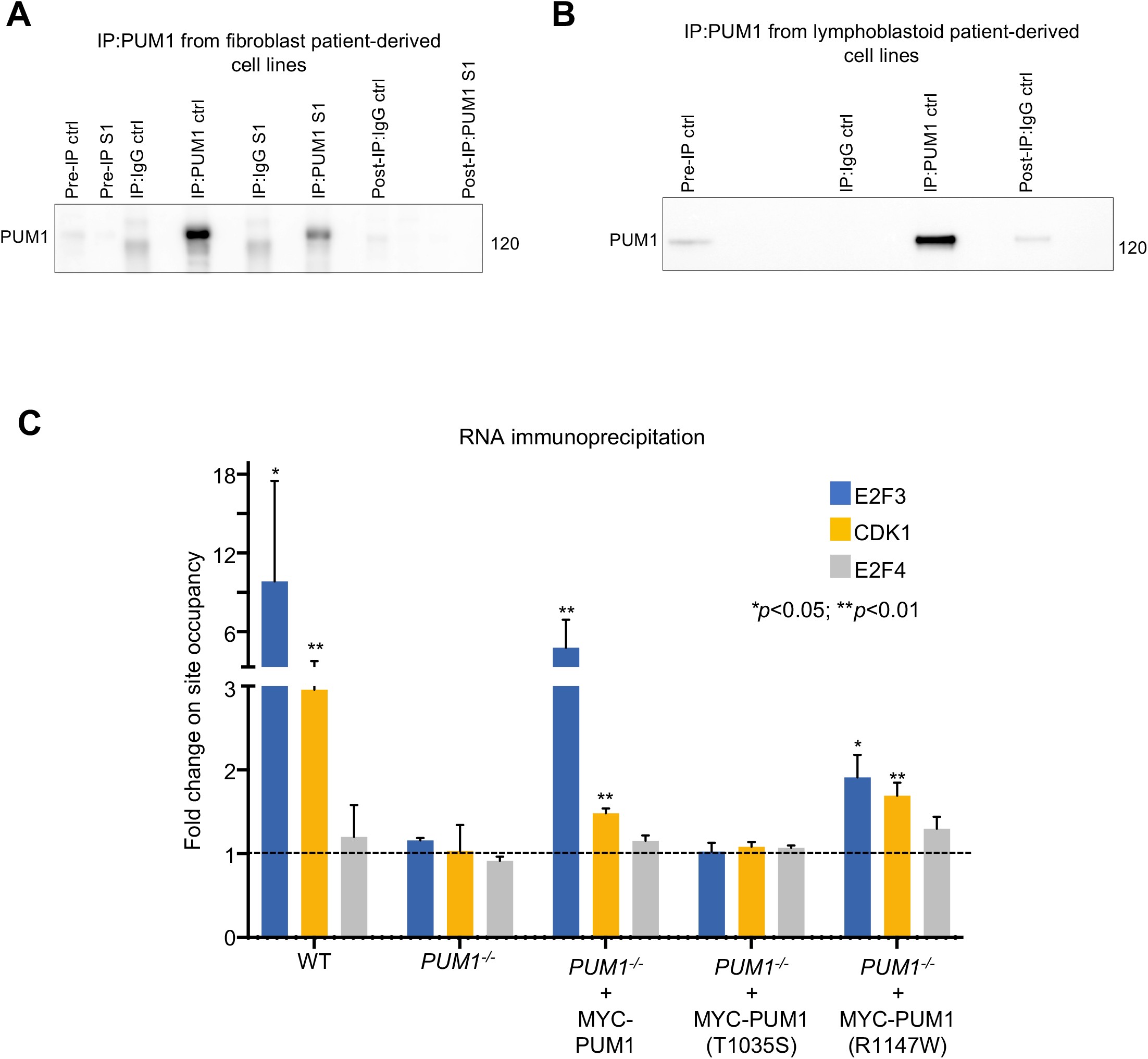
PUM1 R1147W retains RNA-binding activity. (**A**-**B**) Pre-IP, IP, and post-IP against PUM1 and IgG from (**A**) R1147W (infantile-onset SCA7, or PADDAS) fibroblasts and (**B**) T1035S (adult-onset SCA47 or PRCA) lymphoblastoid cells. In both cell lines we were able to pull down 100% of PUM1. Pre-IP represents 1% from the initial protein lysate as a loading control, while 10% of the protein lysate was loaded as post-IP. Molecular weights provided at right in kilodaltons (kDa). (**C**) RNA-immunoprecipitation assay followed by qRT-PCR to show that T1035S is impaired in RNA-binding while R1147W is not. We used *E2F3* and *CDK1* as validated PUM1 targets and E2F4 as a negative control. We transfected WT and PUM1-KO HCT116 cells with 250ng of either Myc-PUM1-WT, Myc-PUM1-R1147W, or Myc-PUM1-T105S. RIP was performed in triplicate.

**Figure S13.**
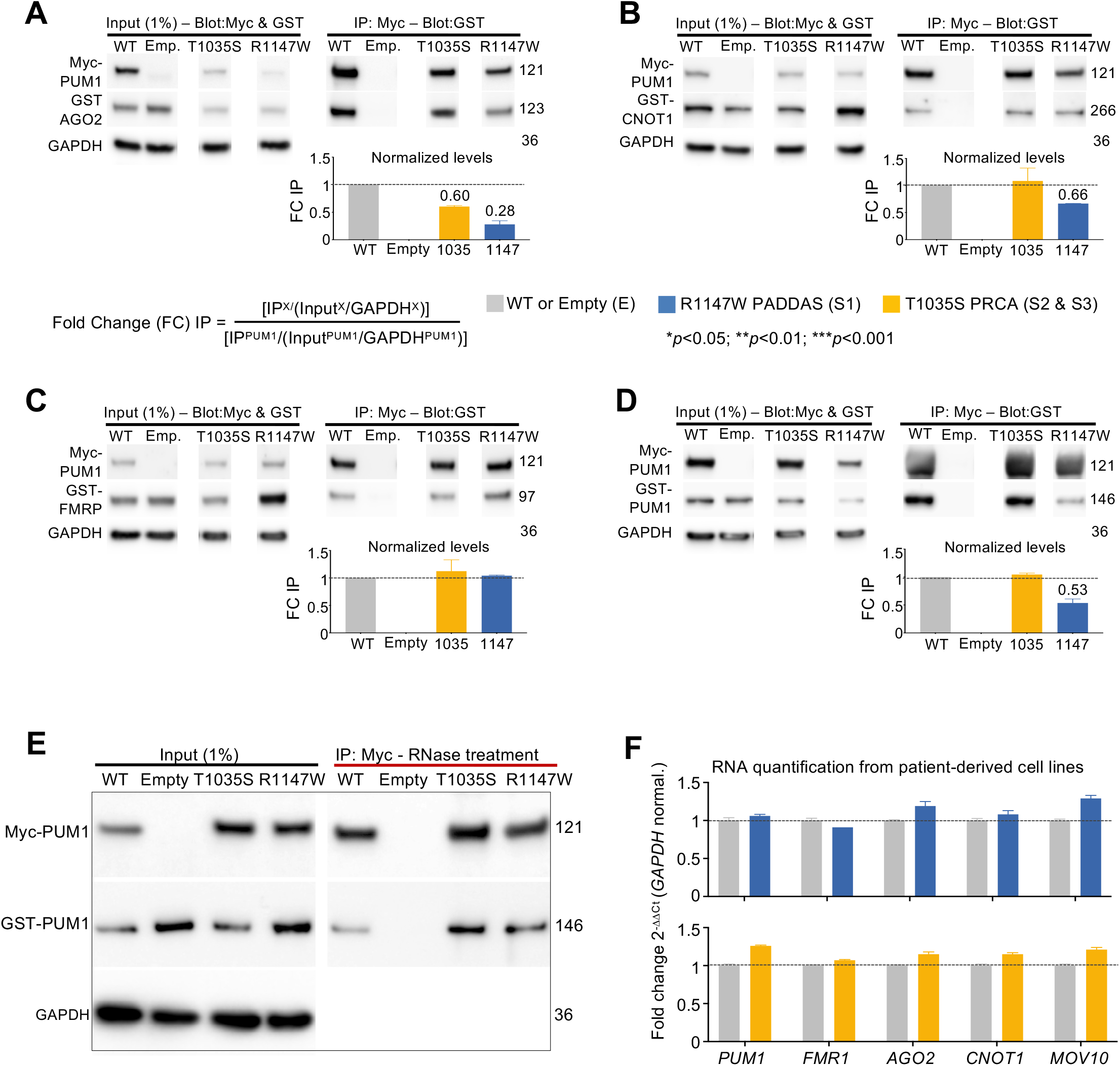
PUM1 validation experiments in patient-derived cells. (**A**-**D**) Representative western blots and relative IP quantification (*bar graphs*) of IP against Myc-PUM1-WT, Myc-PUM1-T1035S (PRCA), or Myc-PUM1-R1147W (PADDAS) followed by immunoblotting for: (**A**) GST-AGO2, (**B**) GST-CNOT1, (**C**) GST-FMRP, and (**D**) GST-PUM1-WT. Myc- and GST-tagged proteins were co-transfected in HEK293T cells in equal quantities (250ng each). The molecular weights were expressed in kilodaltons (kDa). The amount of protein pulled down compared to IP-PUM1 was quantified as [IP^X^/(Input^X^/GAPDH^X^)]/ [IP^PUM1^/(Input^PUM1^/ GAPDH^PUM1^)], where X is the protein of interest. (**E**) Representative western blots of IP with RNase treatment against Myc-PUM1-WT, Myc-PUM1-T1035S (PRCA), and Myc-PUM1-R1147W (PADDAS) followed by immunoblotting to test binding between PUM1 proteins without the RNA. The numbers on the right are the respective molecular weights expressed in kilodaltons (kDa). All the IPs were repeated at least three times. (**F**) mRNA quantification for all of the immunoblotted proteins in Figure 6C in PADDAS and PRCA patient-derived cell lines compared to their respective age-, sex-, and cell-type-matched controls. In all the experiments data represent mean ± SEM. P values were calculated by two-tailed Student’s t test. \**p* < 0.05, \*\**p* < 0.01, \*\*\**p* < 0.001.

**Figure S14.**
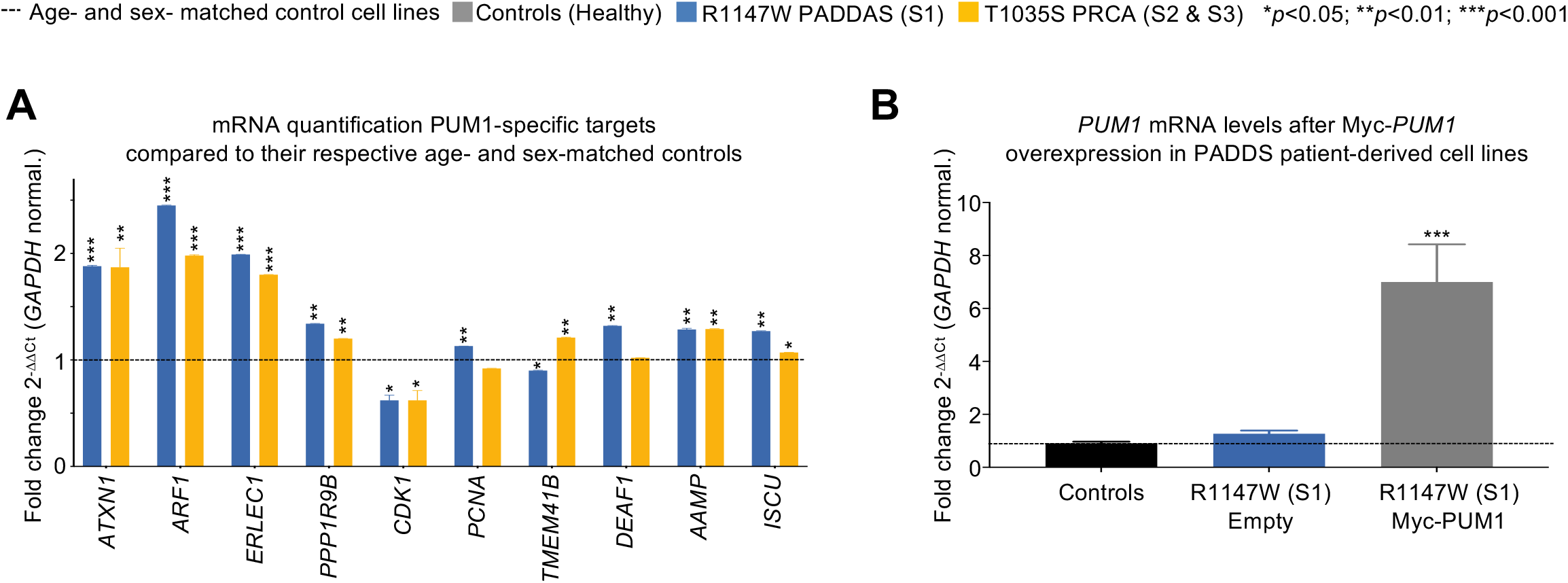
mRNA quantification of PUM1-specific targets. (**A**) qPCR analysis of validated PUM1-specific targets from PADDAS (R1147W) patient-derived fibroblasts (blue bars) compared to three age- and sex-matched fibroblast control cell lines, and PRCA (T1035S) patient-derived lymphoblastoid cell lines (orange bars) compared to three age- and sex-matched lymphoblastoid control cell lines. Only genes expressed in both fibroblasts and lymphoblasts are represented here, for a total of 10 genes. (**B**) mRNA quantification of PUM1 from PADDAS patient-derived fibroblasts transfected with empty and Myc-PUM1-WT vectors, compared to three age- and sex-matched fibroblast control cell lines. All data were normalized to *GAPDH* mRNA levels and experiments performed at least three times. Data represent mean ± SEM. P values were calculated by two-tailed Student’s t test. \**p* < 0.05, \*\**p* < 0.01, \*\*\**p* < 0.001.

## Materials and Methods

### Immunoprecipitation (IP) experiments using mouse brain tissue

Mouse brain tissues were gathered from an equal number of 10-week-old male and female mice. For whole brain experiments, we combined and homogenized two 10-week-old wild-type mouse brains per sample (1 female and 1 male), aliquoting half of each sample for IP against either Pum1 or IgG, then performed six biological replicates (12 mice total) for each LC-MS/MS experiment against IP-Pum1 and IP-IgG. For experiments on the hippocampus, cerebellum, and cortex, we needed much larger numbers of mice: we combined cerebellar and cortical tissues from eight wild-type mice (4 male and 4 female) and performed the experiment in triplicate (total of 24 mice), while for hippocampus we combined tissues from ten wild-type mice (5 female and 5 male) for three experiments (a total of 30 mice).

Samples were processed with a dounce homogenizer using a lysis buffer consisting of 200mM NaCl_2_, 100mM NaPO_4_, 20mM Hepes pH 7.4, 1% Triton X (which should disrupt all but the strongest protein-protein interactions) and complemented by 1X of Xpert Protease and 1X of Phosphatase Inhibitor Cocktail Solutions (GenDepot, #P3100-100, #P3200-020). Following homogenization, the samples were placed on ice for 15 minutes then centrifuged at 14,800 rpm at 4°C for 25 minutes to remove the debris from the supernatant. The supernatant was then moved to 1.5 ml tubes (Beckman microfuge tube #357448) and spun down in a Beckman ultra-centrifuge (Optima Max XP) at 4°C for 25 minutes at 44,000 rpm. 10% of the protein lysate was stored as input and only 1% was loaded for western blot. The protein extract was later divided into two aliquots, one for IP against the protein of interest (antibodies listed below) and the other for IP against IgG, and was then incubated with 30 μL of Dynabeads^TM^ Protein G (Invitrogen, #10004D) and 5 μg of antibody overnight at 4°C on a rotisserie tube rotator. The next day, the beads were washed four times with the same lysis buffer used for IP and resuspended in 40μL of elution buffer (consisting of lysis buffer, NuPAGE 10X Reducing Agent [Invitrogen, #NP0009], NuPAGE LDS sample buffer at 1X final concentration [Invitrogen, #NP0007]) and boiled at 95°C for 10 minutes before the samples were loaded in the NuPAGE 4%–12% Bis-Tris Gels (Invitrogen, #NP0335BOX & #NP0336BOX) for further resolution and western blot analysis.

For the IP with RNase treatment, the beads were resuspended in 400 μL of lysis buffer after the three final washes and divided into two separate 1.5 ml tubes of 200 μL each. To establish the dose required to remove all RNA, we tested different amounts of RNase I (Invitrogen, #EN0602) and found that 4 μL was enough to render RNA undetectable both by denaturing gel and cDNA amplification. This sample and the negative control (i.e., one without RNase treatment) were incubated at 37°C for 15 min on a rotisserie tube rotator. After incubation, all the samples were washed one last time with 500 μL of lysis buffer and then eluted in 20 μL of elution buffer. We used the same protocol for all the IP processed by LC-MS/MS.

The antibodies used for IP were: goat α-PUM1 (Bethyl Laboratories, #A300-201A), rabbit α-PUM2 (Bethyl Laboratories, #A300-202A), rabbit α-FMRP (Abcam Cambridge, #ab17722), rabbit α-AGO2 (Abcam Cambridge, #ab32381), rabbit α-NeuN (Thermo Fisher Scientific, #PA5-37407), rabbit α-CNOT1 (Cell Signaling Technology, #44613), rabbit α-MOV10 (Bethyl Laboratories, #A301-571A), and rabbit α-ANAPC1 (Bethyl Laboratories, #A301-653A).

Please note that *in vivo* IPs from brain lysates present certain challenges that are not encountered *in vitro*. Whereas the total lysate from cells is usually 200μl-300μl, the brain lysate is made in a large volume, usually 1.5 to 2.4 ml, depending on the size of the brain or brain region. This means that in a normal western blot that accommodates 30-40μl total volume, including reducing buffer and loading blue, we cannot load more than 1%-3% from the total brain lysate as input. Therefore, when we pull down a protein of interest (Pum1) and immunoblot for the same protein compared to a standard input (loading the entire IP in one gel), the resulting IP band will be much darker than the input. We then need to expose the Input from the same membrane much longer to visualize it—this is common practice when working with *in vivo* tissues (De Maio *et al*, 2018; Di Grazia *et al*, 2021; Jin *et al*, 2020; Lee *et al*, 2020; Lee *et al*, 2008; Rousseaux *et al*, 2018).

### Immunoprecipitation experiments from HEK293T and patient-derived cell lines

HEK293T cells and patient-derived fibroblasts or lymphoblastoid cells were lysed by pipetting up and down with a 1000μl tip in a lysis buffer consisting of 200mM NaCl_2_, 100mM NaPO_4_, 20mM Hepes pH 7.4, 1% Triton X and complemented by 1X of Xpert Protease and 1X of Phosphatase Inhibitor Cocktail (GenDepot, #P3100-100, #P3200-020). The rest of the protocol is the same as described above for mouse brain tissue, except that we used 2.5 μg of primary antibody for IP.

### Co-Immunoprecipitation in-gel digestion for LC-MS/MS

Immunoprecipitated samples were separated on NuPAGE 4-12% gradient SDS-PAGE (Invitrogen, #NP0335BOX & #NP0336BOX) and stained with SimplyBlue (Invitrogen, #LC6060). Protein gel slices were excised and *in-gel* digestion performed as previously described (Shevchenko *et al*, 2006), with minor modifications. Gel slices were washed with 1:1 Acetonitrile and 100mM ammonium bicarbonate for 30 min then dehydrated with 100% acetonitrile for 10 min until shrunk. The excess acetonitrile was then removed and the slices dried in a speed-vacuum at room temperature for 10 minutes. Gel slices were reduced with 5 mM DTT for 30 min at 56°C in an air thermostat, cooled down to room temperature, and alkylated with 11 mM IAA for 30 min with no light. Gel slices were then washed with 100 mM of ammonium bicarbonate and 100% acetonitrile for 10 min each. Excess acetonitrile was removed and dried in a speed-vacuum for 10 min at room temperature and the gel slices were re-hydrated in a solution of 25 ng/μl trypsin in 50 mM ammonium bicarbonate for 30 min on ice and digested overnight at 37 °C in an air thermostat. Digested peptides were collected and further extracted from gel slices in extraction buffer (1:2 ratio by volume of 5% formic acid: acetonitrile) at high speed, shaking in an air thermostat. The supernatants from both extractions were combined and dried in a speed-vacuum. Peptides were dissolved in 3% acetonitrile/0.1% formic acid.

### Liquid chromatography with tandem mass spectrometry (LC-MS/MS)

The Thermo Scientific Orbitrap Fusion Tribrid mass spectrometer was used for peptide tandem mass spectroscopy (MS/MS). Desalted peptides were injected in an EASY-Spray^TM^ PepMap^TM^ RSLC C18 50cm X 75cm ID column (Thermo Scientific) connected to the Orbitrap Fusion^TM^ Tribrid^TM^. Peptide elution and separation were achieved at a non-linear flow rate of 250 nl/min using a gradient of 5%-30% of buffer B (0.1% (v/v) formic acid, 100% acetonitrile) for 110 minutes, maintaining the temperature of the column at 50 °C during the entire experiment. Survey scans of peptide precursors are performed from 400 to 1500 *m/z* at 120K full width at half maximum (FWHM) resolution (at 200 *m/z*) with a 2 x 10^5^ ion count target and a maximum injection time of 50 ms. The instrument was set to run in top speed mode with 3-second cycles for the survey and the MS/MS scans. After a survey scan, MS/MS was performed on the most abundant precursors, i.e., those ions that had a charge state between 2 and 6, and an intensity of at least 5000, by isolating them in the quadrupole at 1.6 Th. We used collision-induced dissociation (CID) with 35% collision energy and detected the resulting fragments with the rapid scan rate in the ion trap. The automatic gain control (AGC) target for MS/MS was set to 1 x 10^4^ and the maximum injection time was limited to 35ms. The dynamic exclusion was set to 45s with a 10ppm mass tolerance around the precursor and its isotopes. Monoisotopic precursor selection was enabled.

### LC-MS/MS data analysis

Raw mass spectrometric data were analyzed using the MaxQuant environment v.1.6.1.0 (Cox & Mann, 2008) and Andromeda for database searches (Cox *et al*, 2011) at default settings with a few modifications. The default was used for first search tolerance and main search tolerance (20 ppm and 6 ppm, respectively). MaxQuant was set up to search with the reference mouse proteome database downloaded from Uniprot (https://www.uniprot.org/proteomes/UP000000589). MaxQuant searched for trypsin digestion with up to 2 missed cleavages. Peptide, site and protein false discovery rates (FDR) were all set to 1% with a minimum of 1 peptide needed for identification; label-free quantitation (LFQ) was performed with a minimum ratio count of 1. The following modifications were used for protein quantification: oxidation of methionine (M), acetylation of the protein N-terminus, and deamination for asparagine or glutamine (NQ). Results obtained from MaxQuant were further analyzed using the Perseus statistical package (Tyanova *et al*, 2016) that is part of the MaxQuant distribution. Protein identifications were filtered for common contaminants. Proteins were considered for quantification only if they were found in at least two replicate groups. Significant alterations in protein abundance were determined by ANOVA with a threshold for significance of *P* < 0.05 (permutation-based FDR correction). Pum1 protein interactors were later considered if they were found in at least five out of six LC-MS/MS experiments for whole brain and in at least two out of three experiments for each respective brain region with a fold-change of >1.5 between LFQ-PUM1-WT and LFQ-IgG-WT (see **Table S1**).

### Protein-protein interaction map

The protein-protein interaction map for the whole brain (Figure 1) was generated by Cytoscape (https://cytoscape.org/) (Otasek *et al*, 2019) and interactions or functional relationships (clusters) were inferred from Corum (Giurgiu *et al*, 2019) and the Human Protein Atlas (Thul *et al*, 2017) by g:GOSt, which is a package of g:Profiler (https://biit.cs.ut.ee/gprofiler/gost) (Raudvere *et al*., 2019). The brain region-specific map (Figure 2A) was generated by Cytoscape (Shannon *et al*, 2003).

### Quantitative proteomics from mouse brain regions

#### Tissue processing for LC-MS/MS measurements

Frozen brain tissues were weighted and 25-50mg (dry weight) per sample were cryopulverized and lysed in Urea Buffer (8M Urea, 75mM NaCl, 50mM Tris/HCl pH 8.0, 1mM EDTA) in a final 1:10 (mg/µL) tissue:buffer ratio. Protein concentration was determined by BCA assay (Pierce). 40μg of total protein per sample were processed further. Disulfide bonds were reduced with 5mM dithiothreitol and cysteines were subsequently alkylated with 10mM iodoacetamide. Samples were diluted 1:4 with 50mM Tris/HCl (pH 8.0) and sequencing grade modified trypsin (Promega) was added in an enzyme-to-substrate ratio of 1:50. After 16h of digestion, samples were acidified with 1% formic acid (final concentration). Tryptic peptides were desalted on C18 StageTips according to (Rappsilber *et al*, 2007) and evaporated to dryness in a vacuum concentrator. Dried peptides were then reconstituted in 3% ACN / 0.2% Formic acid to a final concentration of 0.5 µg/µL.

#### LC-MS/MS analysis on a Q-Exactive HF

About 1 μg of total peptides were analyzed on a Waters M-Class UPLC using a 25cm Thermo Scientific PepMap RSLC C18 2 µm 25cm column coupled to a benchtop Thermo Fisher Scientific Orbitrap Q Exactive HF mass spectrometer. Peptides were separated at a flow rate of 400 nL/min with a 160 min gradient, including sample loading and column equilibration times. Data was acquired in data independent mode using Xcalibur software. MS1 Spectra were measured with a resolution of 120,000, an AGC target of 5e6 and a mass range from 300 to 1800m/z. MS2 spectra were measured in 47 segment windows per MS1, each with an isolation window width of 32 m/z (0.5 m/z overlap with the neighboring window), a resolution of 30,000, an AGC target of 3e6, and a stepped collision energy of 22.5, 25, 27.5.

All raw data were analyzed with SpectroNaut software version 15.6.211220 (Biognosys) using a directDIA method based on a UniProt mouse database (release 2014_07, Mus musculus), performed with the “BGS factory settings” including the following parameters: Oxidation of methionine and protein N-terminal acetylation as variable modifications; carbamidomethylation as fixed modification; Trypsin/P as the digestion enzyme; For identification, we applied a maximum FDR of 1% separately on protein and peptide level. “Cross run normalization” was activated. This gave intensity values for a total of 5699 protein groups across all samples and replicates. “PG.Quantity” (normalized across samples) values were used for all subsequent analyses.

#### Protein purification of recombinant proteins

Recombinant FMRP, PUM2, and PUM1 were purified from *Escherichia coli* BL21-Codon Plus (DE3)-RIL cells. Both FMRP and PUM2 were expressed with an N-terminal GST tag and isolated from the cell lysate using Glutathione Sepharose 4B (GS4B) resin. The proteins were further subjected to size exclusion chromatography using 20mM HEPES pH 8.0, 200mM NaCl, and 1mM DTT as the buffer. MBP-SNAP-PUM1-His_6_ was expressed and purified similar to the previously described protocol (Elguindy & Mendell, 2021). After nickel affinity chromatography, MBP-SNAP-PUM1-His_6_ was buffer exchanged into 20mM HEPES pH 8.0, 200mM NaCl, and 1mM DTT using PD-10 desalting columns.

#### GST pull-down assay

500mM of purified GST-PUM2 or GST-FMRP were incubated with 1μM of HIS-PUM1 with 30μl of Glutathione Sepharose 4B GST-tagged protein purification resin (Cytiva, #17075601) in a final volume of binding buffer (20mM HEPES pH 8.0, 200mM NaCl, 0.5% Tween-20) for 1 hour at 4°C on the rocker. Resins were washed 3 times in binding buffer and centrifuged at 4000g at 4°C; resins were resuspended in 30μl of elution buffer—which consisted of binding buffer, NuPAGE 10X Reducing Agent (Invitrogen, #NP0009), and NuPAGE LDS sample buffer at 1X final concentration (Invitrogen, #NP0007)—and boiled at 95°C for 10 minutes before the samples were loaded in the NuPAGE 4%–12% Bis-Tris Gels (Invitrogen, #NP0335BOX & #NP0336BOX) for further resolution and western blot analysis. 500mM of purified GST-PUM2, GST-FMRP and HIS-PUM1 were used as input controls; GST-PUM2 and GST-FMRP were visualized using rabbit α-GST antibody (1:1000 [Cell Signaling Technologies, #2625]), whereas for HIS-PUM1 we used rabbit α-HIS Tag antibody (1:3000, [Cell Signaling Technologies, #2365]).

#### Protein quantification and western blot analysis

Patient-derived lymphoblastoid, fibroblast cell lines, and control cell lines were collected at 6 × 10^6^ cell confluence and processed for protein extraction. For mouse tissues, we processed either half of the whole brain (the other half was processed for RNA extraction, see below) or the entire hippocampus, cortex, or cerebellum for protein extraction. Mouse tissues or cell pellets were subsequently lysed with modified RIPA buffer consisting of 25 mM Tris-HCL, pH 7.6, 150 mM NaCl, 1.0% Tween 20, 1.0% sodium deoxycholate, 0.1% SDS, completed with 1X Xpert Protease and 1X Phosphatase Inhibitor Cocktail Solutions (GenDepot, #P3100-100 & #P3200-020). Cells were lysed by pipetting them up and down with a p1000 tip and then placed on ice for 20 min followed by centrifugation at 14,800 rpm at 4°C for 25 minutes. Mouse brain tissues were pipetted up and down by syringe needles—starting from an 18G 1½” (Becton Dickson, #305196), moving to 21G 1½” (Becton Dickson, #305167) and finally to a 26G 1½” (Becton Dickson, #305111) needle—until the lysate passed through the needle smoothly. Proteins were quantified by Pierce BCA Protein Assay Kit (Thermo Scientific, # PI23225) and their absorbance measured by NanoDrop OneC (Thermo Scientific). Proteins were resolved by high resolution NuPAGE 4%–12% Bis-Tris Gel (Invitrogen, #NP0335BOX & #NP0336BOX) according to the manufacturer’s instructions. All the blots were acquired on the G:BOX Chemi XX9 machine (Syngene; Frederick, MD) using GeneSys software 1.6.5.0. Gel exposures were determined by the software.

Antibodies used for western blot experiments were: goat α-PUM1 [1:2500, (Bethyl Laboratories, #A300-201A)], rabbit α-PUM1 [1:2000, (Abcam Cambridge, #ab92545)], rabbit α-PUM2 [1:2000, (Bethyl Laboratories, # A300-202A)], rabbit α-FMRP [1:1000, (Abcam Cambridge, #ab17722)], rabbit α-AGO2 [1:1000, (Abcam Cambridge, #ab32381)], rabbit α-NeuN (Rbfox3) [1:1000, (Thermo Scientific, #PA5-37407)], rabbit α-CNOT1 [1:1000, (Cell Signaling Technology, #44613)], rabbit α-MOV10 [1:2000, (Bethyl Laboratories, #A301-571A)], and mouse α-GAPDH [1:10000, (Millipore, #CB1001)].

#### HEK293T cell culture and maintenance

Human embryonic kidney immortalized 293T (HEK293T) cells were grown in DMEM (GenDepot, #CM002-320) supplemented with 10% of heat-inactivated fetal bovine serum (FBS [GenDepot, #F0901-050) and 1% penicillin/streptomycin (GenDepot, #CA005-010). All cells were incubated at 37 °C in a humidified chamber supplemented with 5% CO_2_. HEK293T cells were later processed according to the needs of specific experiments (described below).

#### RNA extraction and quantitative real-time PCR (qPCR)

Human fibroblast, lymphoblastoid, and respective control cell lines were harvested at 6 × 10^6^ confluence prior to RNA extraction. For mouse tissues, half of the whole brain (the other half was processed for protein extraction, see above) or the entire hippocampus, cortex, or cerebellum were processed for RNA extraction. The RNA was collected for both human cells, mouse brain and brain region tissues using the miRNeasy kit (QIAGEN, # 217004) according to the manufacturer’s instructions. RNA was quantified using NanoDrop OneC (Thermo Fisher Scientific). cDNA was synthesized using Quantitect Reverse Transcription kit (QIAGEN, # 205313) starting from 1 μg of RNA. Quantitative RT-polymerase chain reaction (qRT-PCR) experiments were performed using the CFX96 Touch Real-Time PCR Detection System (Bio-Rad Laboratories, Hercules) with PowerUP SYBR Green Master Mix (Applied Biosystems, #A25743). Real-time PCR runs were analyzed using the comparative C_T_ method normalized against the housekeeping human gene *GAPDH* or mouse *Gapdh*, depending on the experiment (Vandesompele *et al*, 2002).

#### RNA Immunoprecipitation (RIP)

HCT116 wild-type and PUM1-KO were obtained from (Lee *et al*, 2016). HCT116-PUM1-KO cells were grown in McCoy’s 5A Media (Merck, # M4892) supplemented with 10% of heat-inactivated fetal bovine serum (FBS [GenDepot, #F0901-050]) and 1% penicillin/streptomycin (GenDepot, #CA005-010) for 16 hours in a 10-cm plate (80% confluency) and transfected with 6μg of Myc-PUM1-WT, Myc-PUM1-R1147W, and Myc-PUM1-T1035S. After 48 hours cells were collected and lysed in Polysome Lysis Buffer (PLB) (10mM HEPES-KOH pH 7, 100mM KCl, 5mM MgCl_2_, 25 mM EDTA, 0.5% NP40, 2mM Dithiothreitol, 0.2 mg/ml Heparin) supplemented with 50U/ml of RNAse OUT^TM^ recombinant ribonuclease inhibitor (Thermo Fischer Scientific, # 10777019), 50U/ml of SuperaseIN^TM^ RNAse inhibitor (Thermo Fischer Scientific, #AM2694) and 1X of Xpert Protease Inhibitor Cocktail Solutions (GenDepot, #P3100-100) for 20 minutes in ice; cells were centrifuged at maximum speed at 4°C to remove cellular debris. 5μg of goat α-PUM1 (Bethyl Laboratories, #A300-201A) or α-Goat-IgG with 50μl of Dynabeads^TM^ Protein G (Invitrogen, #10004D) were incubated overnight at 4°C on a rotisserie tube rotator; 100μl of lysate were removed to serve as input. 3mg of lysate with 10μl of Dynabeads^TM^ Protein G (Invitrogen, #10004D) were precleared for 30 minutes at 4°C on a rotisserie tube rotator; the pre-cleared lysate was incubated with antibody-conjugated beads overnight at 4°C on a rotisserie tube rotator. Beads were washed 3 times with NT2 buffer (50mM of Tris-HCl, 300mM NaCl, 1mM EDTA, 1% NP40, 0.1% SDS, 0.5% Na-deoxycholate supplemented with 1X of Xpert Protease Inhibitor Cocktail Solutions). Beads and input were resuspended in 100μl of NT2 buffer supplemented with 80U of RNAse OUT and 30μg of proteinase K and incubated for 30 minutes at 50°C to eliminate the proteins. RNA was extracted from the immunoprecipitated samples and their input; 200μl of RLT buffer was added using RNeasy Mini Kit (Qiagen, #74104) according to the manufacturer’s instruction. cDNA was synthetized using SuperScript^TM^ II Reverse Transcripatse (Thermo Fisher Scientific, #18064014) according to the manufacturer’s instructions. Quantitative RT-polymerase chain reaction (qRT-PCR) experiments were performed using the CFX96 Touch Real-Time PCR Detection System (Bio-Rad Laboratories, Hercules) with PowerUP SYBR Green Master Mix (Applied Biosystems, #A25743). The oligos used for qPCR are listed in **Table S7**.

#### MicroRNA library construction and sequencing

Library preparation and microRNA sequencing were performed by LC Sciences (Houston, TX) according to the following criteria. Total RNA was extracted from cerebellum of WT and *Pum1^-/-^*male and female at 10 weeks of age using the miRNeasy kit (QIAGEN, # 217004) according to the manufacturer’s instructions. The total RNA quality and quantity were assessed with Bioanalyzer 2100 (Agilent Technologies, Santa Clara) with RIN number > 7.0. Approximately 1 µg of total RNA were used to prepare the small RNA library according to the protocol of TruSeq Small RNA Sample Prep Kits (Illumina, San Diego). Then the single-end sequencing 50bp was performed on an Illumina Hiseq 2500 at LC Sciences (Hangzhou, China) following the vendor’s recommended protocol.

##### Labelling scheme

| Sample | Gender | Genotype |
| --- | --- | --- |
| LCS7846 GV KO1 F | Female | KO |
| LCS7846 GV KO1 M | Male | KO |
| LCS7846 GV KO2 M | Male | KO |
| LCS7846 GV KO3 F | Female | KO |
| LCS7846 GV KO3 M | Male | KO |
| LCS7846 GV KO4 F | Female | KO |
| LCS7846 GV WT1 M | Male | WT |
| LCS7846 GV WT2 F | Female | WT |
| LCS7846 GV WT2 M | Male | WT |
| LCS7846 GV WT3 F | Female | WT |
| LCS7846 GV WT3 M | Male | WT |

#### MicroRNA sequencing bioinformatic analysis

Raw reads were subjected to an in-house program, ACGT101-miR (LC Sciences, Houston), to remove adapter dimers, junk, common RNA families (rRNA, tRNA, snRNA, snoRNA), and repeats. Subsequently, unique sequences of 18–26 nucleotides in length were mapped to specific species precursors in miRBase 22.0 (http://www.mirbase.org/) by BLAST search to identify known miRNAs and novel 3p-and 5p-derived miRNAs. Length variation at both 3’ and 5’ ends and one mismatch inside of the sequence were allowed in the alignment. The unique sequences mapping to specific species of mature miRNAs in hairpin arms were identified as known miRNAs. The unique sequences mapping to the other arm of known specific species precursor hairpins opposite the annotated mature miRNA-containing arm were considered to be novel 5p- or 3p-derived miRNA candidates. The remaining sequences were mapped to other selected species precursors (with the exclusion of specific species) in miRBase 22.0 by BLAST search, and the mapped pre-miRNAs were further BLASTed against the specific species genomes to determine their genomic locations. The last two were also defined as known miRNAs. The unmapped sequences were BLASTed against the specific genomes, and the hairpin RNA structures containing sequences were predicted from the flank 80 nt sequences using RNAfold software (http://rna.tbi.univie.ac.at/cgi-bin/RNAWebSuite/RNAfold.cgi). The criteria for secondary structure prediction were: (1) number of nucleotides in one bulge in stem (≤12), (2) number of base pairs in the stem region of the predicted hairpin (≥16), (3) cutoff of free energy (kCal/mol ≤-15), (4) length of hairpin (up and down stems + terminal loop ≥50), (5) length of hairpin loop (≤20), (6) number of nucleotides in one bulge in mature region (≤8), (7) number of biased errors in one bulge in mature region (≤4), (8) number of biased bulges in mature region (≤2), (9) number of errors in mature region (≤7), (10) number of base pairs in the mature region of the predicted hairpin (≥12), (11) percent of mature region in stem (≥80).

#### Gene Set Enrichment Analysis (GSEA)

GSEA was performed as previously described (Subramanian *et al*., 2005). The cumulative distribution function was conducted by performing 1000 random gene-set membership assignments. A nominal p-value < 0.01 and an FDR < 0.25 were used to assess the significance of the enrichment score (ES). HITS-CLIP data, and the respective rank, were obtained from the literature and were initially acquired as follows: Pum1 and Pum2 from neonatal murine brains (Zhang *et al*., 2017), Fmrp from cerebellum, cortex, and hippocampus together (Maurin *et al*., 2018), Ago2 from neocortex at embryonic day 13 (Chi *et al*., 2009), Rbfox3 from mouse brain (age not specified) (Weyn-Vanhentenryck *et al*., 2014), Nova from mouse brain (age not specified) (Zhang *et al*., 2010), Ptpb2 from neocortex at embryonic day 18.5 (Licatalosi *et al*., 2012), Mbnl2 from hippocampus at 8-12 weeks of age (Charizanis *et al*., 2012), and Apc from mouse brain at embryonic day 14 (Preitner *et al*., 2014).

#### Gene ontology analyses

Gene ontology analyses were performed with David Gene Ontology (GO). For Figure 2B, Figure S11B and C only categories with FDR<0.05 were considered; while for Figure S10 only categories with FDR<0.01 were considered. David GO for the Pum1 interactome in Figure 2B considered the entire interactome as background. For the GO regarding the HITS-CLIP targets shared among Pum1, Pum2, Fmrp, Ago2, and Rbfox3 (Figure S11B and C), we considered the entire set of all targetomes together as background. Regarding the Synaptic (Syn) GO analysis (Figure S10D), brain-expressed genes were used as background (Koopmans *et al*., 2019).

#### Myc and GST cloning procedure with in vitro immunoprecipitation (IP) assays

Human *PUM1* full-length cDNA was amplified by PCR and subcloned in a pRK5 plasmid containing the Myc tag sequence (Addgene, pRK5-Myc-Parkin #17612) at the N-terminal by using SalI (New England Biolabs, # R3138S) and NotI (New England Biolabs, #R0189S) restriction enzymes to replace *Parkin* with *PUM1*. For GST, the human full-length *PUM1* cDNA was, again, subcloned first in the pRK5 plasmid containing the GST tag sequence (Addgene, pRK5-HA GST RagC wt, #19304) at the N-terminal by using SalI and NotI restriction enzymes to replace *RagC* with *PUM1*. Human *FMRP*, *AGO2* and *CNOT1*, full-length cDNA were cloned and contain the GST tag sequence at the N-terminal, as described for GST-*PUM1*.

To introduce the T1035S or R1147W mutations we used the QuikChange II XL Multi Site-Directed Mutagenesis kit (Agilent Technologies, #200521). The primers for the single mutagenesis experiments were designed by QuikChange software (Stratagene, San Diego, https://www.genomics.agilent.com/primerDesignProgram.jsp).

For IP, HEK293T cells were seeded in 6-well plates for 24 h and then transfected with 250 ng of either WT or mutant PUM1 plasmid with one of the interactors using the jetPRIME Transfection Reagent (Polyplus transfection, #55-132) as per the manufacturer’s protocol. pRK5-Myc empty plasmid (no cDNA) was used as a negative control. After 48 h, the cells were collected and processed for immunoprecipitation. Protein lysates were incubated overnight at 4°C with mouse α-Myc antibody (1:400, [Cell Signaling Technologies, #2276]) on a rotisserie tube rotator. The next day, the beads were washed four times with an IP lysis buffer and resuspended in 40 μL elution buffer (lysis buffer, NuPAGE 10X Reducing Agent [Invitrogen, #NP0009], NuPAGE LDS sample buffer 4X [Invitrogen, #NP0007]) and boiled at 95°C for 10 minutes before loading the samples in NuPAGE 4%–12% Bis-Tris Gels (Invitrogen, #NP0335BOX & #NP0336BOX) for further resolution and western blot analysis. Antibodies: mouse α-Myc antibody [1:2000, (Cell Signaling Technologies, #2276)], rabbit α-GST antibody [1:1000, (Cell Signaling Technologies, #2625)], mouse α-GAPDH [1:10000, (Millipore, #CB1001)].

#### Patient-derived cell lines

Primary fibroblasts from the *PUM1* PADDAS patient (9-year-old female) and the age- and sex-matched controls (three different 9-year-old female) were generated as previously described (Gennarino *et al*., 2018). Briefly, cells were isolated from skin biopsies taken from the patient or age-matched controls using standard methodology (Barch and Association of Cytogenetic Technology, 1991) and placed in a transport medium (Ham’s F10, Thermo Scientific, #11550043). The skin specimen was later removed from the transport medium using a sterile technique (in a Class II biohazard cabinet) and transferred to a sterile Petri dish where it was cut into small pieces (< 0.5 mm) using sterile scalpel blades. These pieces were transferred to the lower surface of a 25 cm^2^ culture flask (6-8 pieces per flask) which had been pre-moistened with 1-2 mL of AmnioMAX Complete Medium (Thermo Scientific, #11269016) supplemented with 1% penicillin/streptomycin (GenDepot, #CA005-010). Cell cultures were maintained at 37 °C in a humidified incubator supplemented with 5% CO_2_. When cell growth was observed around the edges of the tissue, usually 3 to 5 days later, 2 to 3 mL of AmnioMAX Complete Medium were added. Once growth was established and the tissue was anchored to the flask, another 8 mL of AmnioMAX Complete Medium was added. Thereafter, the medium was renewed every 3 to 4 days until ready for sub-culturing.

Lymphoblastoid cells from *PUM1* PRCA patients (female 59 and 58 years old, respectively) and the age- and sex-matched controls (three different 58-year-old female) were generated as previously described (Gennarino *et al*., 2018). Briefly, lymphoblastoid suspension cell cultures were grown in RPMI 1640 medium (Invitrogen, #11875093) supplemented with 10% heat-inactivated fetal bovine serum (Atlanta Biological, Flowery Branch, #S11195H) and 1% penicillin/streptomycin (GenDepot, #CA005-010). Cell cultures were maintained at 37°C in a humidified incubator supplemented with 5% CO_2_. Medium was renewed every 2 to 3 days.

#### Fibroblast patient-derived cell lines transfection

Fibroblasts from age- and sex-matched healthy controls and from a female PADDAS patient were seeded at 80% of confluency in 6-well plates (∼150.000 cells/well). The day after, 500ng of pRK5-CMV-Myc-Pum1 or pRK5-CMV-Myc-Empty plasmids were transfected in antibiotic-free DMEM (GenDepot, #CM002-320) using Lipofectamine LTX with Plus Reagent (Thermo Fisher, #15338030) according to the manufacturer’s protocol. After 5 hours we replaced the media with new complete DMEM supplemented with 10% of heat-inactivated fetal bovine serum (FBS [GenDepot, #F0901-050]) and 1% penicillin/streptomycin (GenDepot, #CA005-010). Cells were incubated at 37 °C in a humidified chamber supplemented with 5% CO2 and collected after 72 hours for RNA and protein extraction.

#### Primers

For the qPCR analysis to unambiguously distinguish spliced cDNA from genomic DNA contamination, specific exon primers were designed to amplify across introns of the gene tested. The primers for all genes tested were designed with Primer3 (Koressaar & Remm, 2007; Untergasser *et al*, 2012). Cloning primers were manually designed to amplify the longest spliced gene isoform tested; if there was more than one isoform according to the UCSC Genome Browser (https://genome.ucsc.edu/), we chose the longest. See Table S7 for primer sequences.

#### Ethical statement and mouse strains

All animal procedures were approved by the Institutional Animal Care and Use Committee at Columbia University, New York under the protocol AC-AAAU8490. Mice were maintained on a 12-hr light, 12-hr dark cycle with regular chow and water ad libitum. Pum1 knock-out mice were generated as previously described (Chen *et al*., 2012). C57BL/6J wild-type mice were purchased from Jackson Laboratory and maintained as described above. For brain dissection, mice were anesthetized with isoflurane, and the brain rapidly removed from the skull and lysed in the appropriate buffer according to the experiment (see Materials and Methods Details).

#### Experimental design

For protein and RNA quantification from patient-derived cell lines, we used values from at least six independent experiments with three biological replicates for each experiment. At every stage of the study, the experimenter was blinded to the identity of control and patient-derived cell lines. For example, for the data regarding both human patient-derived cell lines and mice, Experimenter #1 made a list of samples and controls to be tested, and Experimenter #2 randomized this list and re-labeled the tubes; Experimenter #2 was the only person with the key to identify the samples. These samples were then distributed to Experimenter #3 to culture the cells, then to Experimenter #1 to perform western blots and qRT-PCR, and lastly Experimenters #1 and #4 analyzed the data. Only then was the key applied to identify the samples.

For mouse experiments, the experimenters were randomized and blinded as described above. The number of animals used and sex, and the specific statistical tests used, are indicated for each experiment in the figure legends. Sample size was based on previous experience using the same mice (Gennarino *et al*., 2015).

#### Software and statistical analysis

Statistical significance was analyzed using GraphPad Prism 8 (https://www.graphpad.com/scientific-software/prism/) and Excel Software (Microsoft). All data are presented as mean ± SEM. Statistical details for each experiment can be found in the figures and the legends. The range of expression levels in qPCR was determined from at least six independent experiments with three biological replicates by calculating the standard deviation of the **Δ**Ct (Pfaffl, 2001). The range of expression levels in western blots was determined from at least six independent experiments with at least six biological replicates. P values were calculated by Student’s T-test or analysis of variance with Tukey’s post hoc analysis. For the IP in Figure 6A and protein quantification in patient cell lines in Figure 6C, we had only one PADDAS patient, so the repeated experiments were technical replicates rather than biological replicates. We therefore calculated the statistical significance based on these technical replicates in comparison to the three biological replicates (i.e., healthy controls).

#### Study approval

PADDAS and PRCA patient cell lines are the same as those reported previously (Gennarino *et al*., 2018). The consent form for each subject specifically allows for sharing of medical information and physical exam findings; the sharing of cell lines from the PADDAS and PRCA subjects and the controls was approved under the Columbia University Medical Center IRB-AAAS7401 (Y01M00) and the Baylor College of Medicine IRB H-34578.

### Data availability

IP mass spec and quantitative proteomics raw data are available at: “pending”. microRNA sequencing raw data are available at “pending”

## Notes

### Competing Interest Statement

The authors have declared no competing interest.

### Summary of Updates

Figure 3 and 4 revised. Figure 2 revised. Figure 7 revised Added Figure S3 Added Figure S4.

## References

Agarwal V, Bell GW, Nam JW, Bartel DP (2015) Predicting effective microRNA target sites in mammalian mRNAs. Elife 4

Bah MG, Rodriguez D, Cazeneuve C, Mochel F, Devos D, Suppiej A, Roubertie A, Meunier I, Gitiaux C, Curie A et al (2020) Deciphering the natural history of SCA7 in children. Eur J Neurol 27: 2267–2276

Blackinton JG, Keene JD (2014) Post-transcriptional RNA regulons affecting cell cycle and proliferation. Semin Cell Dev Biol 34: 44–54

Blewett NH, Goldstrohm AC (2012) A eukaryotic translation initiation factor 4E-binding protein promotes mRNA decapping and is required for PUF repression. Mol Cell Biol 32: 4181–4194

Bohn JA, Van Etten JL, Schagat TL, Bowman BM, McEachin RC, Freddolino PL, Goldstrohm AC (2018) Identification of diverse target RNAs that are functionally regulated by human Pumilio proteins. Nucleic Acids Res 46: 362–386

Bonnemason-Carrere P, Morice-Picard F, Pennamen P, Arveiler B, Fergelot P, Goizet C, Hellegouarch M, Lacombe D, Plaisant C, Raclet V et al (2019) PADDAS syndrome associated with hair dysplasia caused by a de novo missense variant of PUM1. Am J Med Genet A 179: 1030–1033

Charizanis K, Lee KY, Batra R, Goodwin M, Zhang C, Yuan Y, Shiue L, Cline M, Scotti MM, Xia G et al (2012) Muscleblind-like 2-mediated alternative splicing in the developing brain and dysregulation in myotonic dystrophy. Neuron 75: 437–450

Chartier-Harlin MC, Kachergus J, Roumier C, Mouroux V, Douay X, Lincoln S, Levecque C, Larvor L, Andrieux J, Hulihan M et al (2004) Alpha-synuclein locus duplication as a cause of familial Parkinson’s disease. Lancet 364: 1167–1169

Chen D, Zheng W, Lin A, Uyhazi K, Zhao H, Lin H (2012) Pumilio 1 suppresses multiple activators of p53 to safeguard spermatogenesis. Curr Biol 22: 420–425

Chi SW, Zang JB, Mele A, Darnell RB (2009) Argonaute HITS-CLIP decodes microRNA-mRNA interaction maps. Nature 460: 479–486

Conboy JG (2017) Developmental regulation of RNA processing by Rbfox proteins. Wiley Interdiscip Rev RNA 8

Cox J, Mann M (2008) MaxQuant enables high peptide identification rates, individualized p.p.b.-range mass accuracies and proteome-wide protein quantification. Nat Biotechnol 26: 1367–1372

Cox J, Neuhauser N, Michalski A, Scheltema RA, Olsen JV, Mann M (2011) Andromeda: a peptide search engine integrated into the MaxQuant environment. J Proteome Res 10: 1794–1805

Darnell JC, Richter JD (2012) Cytoplasmic RNA-binding proteins and the control of complex brain function. Cold Spring Harb Perspect Biol 4: a012344

Dassi E (2017) Handshakes and Fights: The Regulatory Interplay of RNA-Binding Proteins. Front Mol Biosci 4: 67

De Maio A, Yalamanchili HK, Adamski CJ, Gennarino VA, Liu Z, Qin J, Jung SY, Richman R, Orr H, Zoghbi HY (2018) RBM17 Interacts with U2SURP and CHERP to Regulate Expression and Splicing of RNA-Processing Proteins. Cell Rep 25: 726–736 e727

Di Grazia A, Marafini I, Pedini G, Di Fusco D, Laudisi F, Dinallo V, Rosina E, Stolfi C, Franze E, Sileri P et al (2021) The Fragile X Mental Retardation Protein Regulates RIPK1 and Colorectal Cancer Resistance to Necroptosis. Cell Mol Gastroenterol Hepatol 11: 639–658

Elguindy MM, Mendell JT (2021) NORAD-induced Pumilio phase separation is required for genome stability. Nature 595: 303–308

Enwerem, III, Elrod ND, Chang CT, Lin A, Ji P, Bohn JA, Levdansky Y, Wagner EJ, Valkov E, Goldstrohm AC (2021) Human Pumilio proteins directly bind the CCR4-NOT deadenylase complex to regulate the transcriptome. RNA 27: 445–464

Friend K, Campbell ZT, Cooke A, Kroll-Conner P, Wickens MP, Kimble J (2012) A conserved PUF-Ago-eEF1A complex attenuates translation elongation. Nat Struct Mol Biol 19: 176–183

Fu J, Peng L, Tao T, Chen Y, Li Z, Li J (2019) Regulatory roles of the miR-200 family in neurodegenerative diseases. Biomed Pharmacother 119: 109409

Gehman LT, Meera P, Stoilov P, Shiue L, O’Brien JE, Meisler MH, Ares M, Jr., Otis TS, Black DL (2012) The splicing regulator Rbfox2 is required for both cerebellar development and mature motor function. Genes Dev 26: 445–460

Genetic Modifiers of Huntington’s Disease C (2015) Identification of Genetic Factors that Modify Clinical Onset of Huntington’s Disease. Cell 162: 516–526

Gennarino VA, D’Angelo G, Dharmalingam G, Fernandez S, Russolillo G, Sanges R, Mutarelli M, Belcastro V, Ballabio A, Verde P et al (2012) Identification of microRNA-regulated gene networks by expression analysis of target genes. Genome Res 22: 1163–1172

Gennarino VA, Palmer EE, McDonell LM, Wang L, Adamski CJ, Koire A, See L, Chen CA, Schaaf CP, Rosenfeld JA et al (2018) A mild PUM1 mutation is associated with adult-onset ataxia, whereas haploinsufficiency causes developmental delay and seizures. Cell 172: 924–936 e911

Gennarino VA, Singh RK, White JJ, De Maio A, Han K, Kim JY, Jafar-Nejad P, di Ronza A, Kang H, Sayegh LS et al (2015) Pumilio1 haploinsufficiency leads to SCA1-like neurodegeneration by increasing wild-type Ataxin1 levels. Cell 160: 1087-1098

Giurgiu M, Reinhard J, Brauner B, Dunger-Kaltenbach I, Fobo G, Frishman G, Montrone C, Ruepp A (2019) CORUM: the comprehensive resource of mammalian protein complexes-2019. Nucleic Acids Res 47: D559-D563

Goldstrohm AC, Hall TMT, McKenney KM (2018) Post-transcriptional Regulatory Functions of Mammalian Pumilio Proteins. Trends Genet 34: 972–990

Hentze MW, Castello A, Schwarzl T, Preiss T (2018) A brave new world of RNA-binding proteins. Nat Rev Mol Cell Biol 19: 327–341

Imaizumi T, Mogami Y, Okamoto N, Yamamoto-Shimojima K, Yamamoto T (2019) De novo 1p35.2 microdeletion including PUM1 identified in a patient with sporadic West syndrome. Congenit Anom (Kyoto) 59: 193–194

Jin Z, Feng H, Liang J, Jing X, Zhao Q, Zhan L, Shen B, Cheng X, Su L, Qiu W (2020) FGFR3 big up tri, open7-9 promotes tumor progression via the phosphorylation and destabilization of ten-eleven translocation-2 in human hepatocellular carcinoma. Cell Death Dis 11: 903

Kacher R, Lejeune FX, Noel S, Cazeneuve C, Brice A, Humbert S, Durr A (2021) Propensity for somatic expansion increases over the course of life in Huntington disease. Elife 10

Kedde M, van Kouwenhove M, Zwart W, Oude Vrielink JA, Elkon R, Agami R (2010) A Pumilio-induced RNA structure switch in p27-3’ UTR controls miR-221 and miR-222 accessibility. Nat Cell Biol 12: 1014–1020

Keene JD (2007a) Biological clocks and the coordination theory of RNA operons and regulons. Cold Spring Harb Symp Quant Biol 72: 157–165

Keene JD (2007b) RNA regulons: coordination of post-transcriptional events. Nat Rev Genet 8: 533–543

Keene JD, Lager PJ (2005) Post-transcriptional operons and regulons co-ordinating gene expression. Chromosome Res 13: 327–337

Kenny PJ, Kim M, Skariah G, Nielsen J, Lannom MC, Ceman S (2020) The FMRP-MOV10 complex: a translational regulatory switch modulated by G-Quadruplexes. Nucleic Acids Res 48: 862–878

Kenny PJ, Zhou H, Kim M, Skariah G, Khetani RS, Drnevich J, Arcila ML, Kosik KS, Ceman S (2014) MOV10 and FMRP regulate AGO2 association with microRNA recognition elements. Cell Rep 9: 1729–1741

Khalil B, Morderer D, Price PL, Liu F, Rossoll W (2018) mRNP assembly, axonal transport, and local translation in neurodegenerative diseases. Brain Res 1693: 75–91

Kim K, Hessl D, Randol JL, Espinal GM, Schneider A, Protic D, Aydin EY, Hagerman RJ, Hagerman PJ (2019) Association between IQ and FMR1 protein (FMRP) across the spectrum of CGG repeat expansions. PLoS One 14: e0226811

Koopmans F, van Nierop P, Andres-Alonso M, Byrnes A, Cijsouw T, Coba MP, Cornelisse LN, Farrell RJ, Goldschmidt HL, Howrigan DP et al (2019) SynGO: An Evidence-Based, Expert-Curated Knowledge Base for the Synapse. Neuron 103: 217–234 e214

Koressaar T, Remm M (2007) Enhancements and modifications of primer design program Primer3. Bioinformatics 23: 1289–1291

Lai KL, Liao YC, Tsai PC, Hsiao CT, Soong BW, Lee YC (2019) Investigating PUM1 mutations in a Taiwanese cohort with cerebellar ataxia. Parkinsonism Relat Disord 66: 220–223

Lee EJ, Seo E, Kim JW, Nam SA, Lee JY, Jun J, Oh S, Park M, Jho EH, Yoo KH et al (2020) TAZ/Wnt-beta-catenin/c-MYC axis regulates cystogenesis in polycystic kidney disease. Proc Natl Acad Sci U S A 117: 29001–29012

Lee S, Kopp F, Chang TC, Sataluri A, Chen B, Sivakumar S, Yu H, Xie Y, Mendell JT (2016) Noncoding RNA NORAD Regulates Genomic Stability by Sequestering PUMILIO Proteins. Cell 164: 69–80

Lee Y, Samaco RC, Gatchel JR, Thaller C, Orr HT, Zoghbi HY (2008) miR-19, miR-101 and miR-130 co-regulate ATXN1 levels to potentially modulate SCA1 pathogenesis. Nat Neurosci 11: 1137-1139

Licatalosi DD, Yano M, Fak JJ, Mele A, Grabinski SE, Zhang C, Darnell RB (2012) Ptbp2 represses adult-specific splicing to regulate the generation of neuronal precursors in the embryonic brain. Genes Dev 26: 1626–1642

Marrero E, Rossi SG, Darr A, Tsoulfas P, Rotundo RL (2011) Translational regulation of acetylcholinesterase by the RNA-binding protein Pumilio-2 at the neuromuscular synapse. J Biol Chem 286: 36492–36499

Mauger O, Lemoine F, Scheiffele P (2016) Targeted Intron Retention and Excision for Rapid Gene Regulation in Response to Neuronal Activity. Neuron 92: 1266–1278

Maurin T, Lebrigand K, Castagnola S, Paquet A, Jarjat M, Popa A, Grossi M, Rage F, Bardoni B (2018) HITS-CLIP in various brain areas reveals new targets and new modalities of RNA binding by fragile X mental retardation protein. Nucleic Acids Res 46: 6344–6355

Miles WO, Tschop K, Herr A, Ji JY, Dyson NJ (2012) Pumilio facilitates miRNA regulation of the E2F3 oncogene. Genes Dev 26: 356–368

Muraro NI, Weston AJ, Gerber AP, Luschnig S, Moffat KG, Baines RA (2008) Pumilio binds para mRNA and requires Nanos and Brat to regulate sodium current in Drosophila motoneurons. J Neurosci 28: 2099–2109

Orr HT, Zoghbi HY (2007) Trinucleotide repeat disorders. Annu Rev Neurosci 30: 575–621

Otasek D, Morris JH, Boucas J, Pico AR, Demchak B (2019) Cytoscape Automation: empowering workflow-based network analysis. Genome Biol 20: 185

Pfaffl MW (2001) A new mathematical model for relative quantification in real-time RT-PCR. Nucleic Acids Res 29: e45

Preitner N, Quan J, Nowakowski DW, Hancock ML, Shi J, Tcherkezian J, Young-Pearse TL, Flanagan JG (2014) APC is an RNA-binding protein, and its interactome provides a link to neural development and microtubule assembly. Cell 158: 368–382

Rappsilber J, Mann M, Ishihama Y (2007) Protocol for micro-purification, enrichment, pre-fractionation and storage of peptides for proteomics using StageTips. Nat Protoc 2: 1896–1906

Raudvere U, Kolberg L, Kuzmin I, Arak T, Adler P, Peterson H, Vilo J (2019) g:Profiler: a web server for functional enrichment analysis and conversions of gene lists (2019 update). Nucleic Acids Res 47: W191–W198

Ravanidis S, Kattan FG, Doxakis E (2018) Unraveling the Pathways to Neuronal Homeostasis and Disease: Mechanistic Insights into the Role of RNA-Binding Proteins and Associated Factors. Int J Mol Sci 19

Rousseaux MWC, Tschumperlin T, Lu HC, Lackey EP, Bondar VV, Wan YW, Tan Q, Adamski CJ, Friedrich J, Twaroski K, et al (2018) ATXN1-CIC Complex Is the Primary Driver of Cerebellar Pathology in Spinocerebellar Ataxia Type 1 through a Gain-of-Function Mechanism. Neuron 97: 1235–1243 e1235

Rovelet-Lecrux A, Hannequin D, Raux G, Le Meur N, Laquerriere A, Vital A, Dumanchin C, Feuillette S, Brice A, Vercelletto M et al (2006) APP locus duplication causes autosomal dominant early-onset Alzheimer disease with cerebral amyloid angiopathy. Nat Genet 38: 24–26

Salcedo-Arellano MJ, Dufour B, McLennan Y, Martinez-Cerdeno V, Hagerman R (2020) Fragile X syndrome and associated disorders: Clinical aspects and pathology. Neurobiol Dis 136: 104740

Shah S, Molinaro G, Liu B, Wang R, Huber KM, Richter JD (2020) FMRP Control of Ribosome Translocation Promotes Chromatin Modifications and Alternative Splicing of Neuronal Genes Linked to Autism. Cell Rep 30: 4459–4472 e4456

Shannon P, Markiel A, Ozier O, Baliga NS, Wang JT, Ramage D, Amin N, Schwikowski B, Ideker T (2003) Cytoscape: a software environment for integrated models of biomolecular interaction networks. Genome Res 13: 2498–2504

Shevchenko A, Tomas H, Havlis J, Olsen JV, Mann M (2006) In-gel digestion for mass spectrometric characterization of proteins and proteomes. Nat Protoc 1: 2856–2860

Singh K, Gaur P, Prasad S (2007) Fragile x mental retardation (Fmr-1) gene expression is down regulated in brain of mice during aging. Mol Biol Rep 34: 173–181

Singh K, Prasad S (2008) Differential expression of Fmr-1 mRNA and FMRP in female mice brain during aging. Mol Biol Rep 35: 677–684

Smidak R, Sialana FJ, Kristofova M, Stojanovic T, Rajcic D, Malikovic J, Feyissa DD, Korz V, Hoeger H, Wackerlig J et al (2017) Reduced Levels of the Synaptic Functional Regulator FMRP in Dentate Gyrus of the Aging Sprague-Dawley Rat. Front Aging Neurosci 9: 384

Subramanian A, Tamayo P, Mootha VK, Mukherjee S, Ebert BL, Gillette MA, Paulovich A, Pomeroy SL, Golub TR, Lander ES et al (2005) Gene set enrichment analysis: a knowledge-based approach for interpreting genome-wide expression profiles. Proc Natl Acad Sci U S A 102: 15545–15550

Takeguchi R, Takahashi S, Kuroda M, Tanaka R, Suzuki N, Tomonoh Y, Ihara Y, Sugiyama N, Itoh M (2020) MeCP2_e2 partially compensates for lack of MeCP2_e1: A male case of Rett syndrome. Mol Genet Genomic Med 8: e1088

Tan Q, Zoghbi HY (2019) Mouse models as a tool for discovering new neurological diseases. Neurobiol Learn Mem 165: 106902

Temme C, Simonelig M, Wahle E (2014) Deadenylation of mRNA by the CCR4-NOT complex in Drosophila: molecular and developmental aspects. Front Genet 5: 143

Tereshchenko A, Magnotta V, Epping E, Mathews K, Espe-Pfeifer P, Martin E, Dawson J, Duan W, Nopoulos P (2019) Brain structure in juvenile-onset Huntington disease. Neurology 92: e1939–e1947

Thul PJ, Akesson L, Wiking M, Mahdessian D, Geladaki A, Ait Blal H, Alm T, Asplund A, Bjork L, Breckels LM et al (2017) A subcellular map of the human proteome. Science 356

Trumbach D, Prakash N (2015) The conserved miR-8/miR-200 microRNA family and their role in invertebrate and vertebrate neurogenesis. Cell Tissue Res 359: 161–177

Tyanova S, Temu T, Sinitcyn P, Carlson A, Hein MY, Geiger T, Mann M, Cox J (2016) The Perseus computational platform for comprehensive analysis of (prote)omics data. Nat Methods 13: 731–740

Untergasser A, Cutcutache I, Koressaar T, Ye J, Faircloth BC, Remm M, Rozen SG (2012) Primer3--new capabilities and interfaces. Nucleic Acids Res 40: e115

Uyhazi KE, Yang Y, Liu N, Qi H, Huang XA, Mak W, Weatherbee SD, de Prisco N, Gennarino VA, Song X et al (2020) Pumilio proteins utilize distinct regulatory mechanisms to achieve complementary functions required for pluripotency and embryogenesis. Proc Natl Acad Sci U S A 117: 7851–7862

Van Etten J, Schagat TL, Hrit J, Weidmann CA, Brumbaugh J, Coon JJ, Goldstrohm AC (2012) Human Pumilio proteins recruit multiple deadenylases to efficiently repress messenger RNAs. J Biol Chem 287: 36370–36383

Vandesompele J, De Preter K, Pattyn F, Poppe B, Van Roy N, De Paepe A, Speleman F (2002) Accurate normalization of real-time quantitative RT-PCR data by geometric averaging of multiple internal control genes. Genome Biol 3

Wamsley B, Jaglin XH, Favuzzi E, Quattrocolo G, Nigro MJ, Yusuf N, Khodadadi-Jamayran A, Rudy B, Fishell G (2018) Rbfox1 Mediates Cell-type-Specific Splicing in Cortical Interneurons. Neuron 100: 846–859 e847

Weidmann CA, Raynard NA, Blewett NH, Van Etten J, Goldstrohm AC (2014) The RNA binding domain of Pumilio antagonizes poly-adenosine binding protein and accelerates deadenylation. RNA 20: 1298–1319

Weyn-Vanhentenryck SM, Mele A, Yan Q, Sun S, Farny N, Zhang Z, Xue C, Herre M, Silver PA, Zhang MQ et al (2014) HITS-CLIP and integrative modeling define the Rbfox splicing-regulatory network linked to brain development and autism. Cell Rep 6: 1139–1152

Zamore PD, Williamson JR, Lehmann R (1997) The Pumilio protein binds RNA through a conserved domain that defines a new class of RNA-binding proteins. RNA 3: 1421–1433

Zhang C, Frias MA, Mele A, Ruggiu M, Eom T, Marney CB, Wang H, Licatalosi DD, Fak JJ, Darnell RB (2010) Integrative modeling defines the Nova splicing-regulatory network and its combinatorial controls. Science 329: 439–443

Zhang M, Chen D, Xia J, Han W, Cui X, Neuenkirchen N, Hermes G, Sestan N, Lin H (2017) Post-transcriptional regulation of mouse neurogenesis by Pumilio proteins. Genes Dev 31: 1354–1369

